# Karyotypic evolution in a diverse primate clade and the role of chromosomal fissions as barriers to gene flow

**DOI:** 10.64898/2026.09.14.751345

**Authors:** Axel Jensen, Athena Syarifa, Julien Pichon, Micaela Strada, Fanny Combe, Bertrand Bed’hom, Loic Ponger, Michèle Gerbault-Seureau, Christophe Escudé, Katerina Guschanski

## Abstract

The classical model of chromosomal speciation is based on the observation that divergent species often differ in chromosome numbers and that heterokaryotypic hybrids show reduced fitness. Furthermore, positive correlations between the rates of karyotype evolution and speciation have been established in several groups of organisms. Yet, the mechanisms that make some lineages more prone to chromosomal fissions and fusions than others, and the causal role of such karyotype changes in speciation, are poorly understood. Here, we focus on the taxonomically and karyotypically diverse group of African monkeys, the guenons (tribe Cercopithecini), consisting of ∼35 species with diploid chromosome numbers ranging from 48 to 72. We generated *de novo* genome assemblies for eight species with different karyotypes, and incorporated publicly available genomes from additionally three guenon species. Using the rhesus macaque as an outgroup, we identified 28 independent chromosomal fissions and 4 fusions in guenons. In line with previous cytogenetic studies, repeat composition around chromosomal breakpoints suggested that most fissions were non-centromeric, requiring the evolution of a new centromere in at least one of the fission products. In contrast to the expectation under strong underdominance, we found a paraphyletic distribution of several chromosomal fissions. Some of these phylogenetically discordant events could have resulted from interspecific hybridization, whereas others are more likely caused by incomplete lineage sorting, suggesting that fissioned and non-fissioned homologs segregated in ancestral guenon lineages for millions of years. Nevertheless, we find evidence for reduced introgression on rearranged chromosomes in an ancient cross-karyotypic gene flow event between *Cercopithecus cephus* and *C. pogonias*. Although this suggests that chromosomal fissions may have constituted barriers to gene flow in the guenon evolution, the signal was inconclusive or absent in two other investigated gene flow events. This study provides important first insights into the remarkable karyotypic diversification of guenons, and its role in generating reproductive barriers.

## Introduction

The emergence of novel species is an intensively studied topic in evolutionary biology. Yet, the mechanisms underlying the evolution of reproductive isolation, a prerequisite for speciation, remain incompletely understood. Chromosomal reorganization was one of the first proposed reproductive isolation mechanism (Sturtevant, 1938; White, 1968), with early models motivated by the observations that closely related species often harbor karyotypic differences, and that heterokaryotypic crosses are often infertile (Bush et al., 1977; White, 1968).

Two main “sub-models” of chromosomal speciation have been proposed (Ayala & Coluzzi, 2005). The ‘suppressed recombination model’ postulates that rearranged parts of the genome will experience reduced recombination in heterozygotes, thus becoming hotspots for accumulation of genomic incompatibilities and alleles involved in local adaptation (Rieseberg 2001). This could facilitate species divergence, as gene flow will be locally reduced between lineages that differ in such rearrangements. The suppressed recombination model predicts that emerging species pairs will be more divergent in and around rearranged regions of the genome, which has indeed been demonstrated in several systems (e.g. Faria et al., 2019; Lowry & Willis, 2010). The ‘hybrid dysfunction model’ of chromosomal speciation, rests solely on the reduced fertility of heterokaryotypic hybrids (White, 1968). Meiosis in heterokaryotypic hybrids may produce unbalanced gametes, and thus reduced fertility, which is suggested to acts as a strong reproductive barrier. While the suppressed recombination model states the importance of smaller scale rearrangements, such as intrachromosomal translocations and inversions, the hybrid dysfunction model is more often attributed to fissions and fusions, i.e., lineages that differ in chromosome numbers.

Importantly, the hybrid dysfunction model of chromosomal speciation suffers from a potentially problematic paradox: Karyotypic changes that cause strong underdominance, that is, heterozygous disadvantage, are unlikely to establish in the first place (Coyne & Orr, 2004). When a chromosomal fission or fusion first occurs, it will be rare and thus more likely to occur in a heterozygous state with the ancestral karyotype. If underdominance is strong, the new karyotype should be rapidly purged by purifying selection. Suggested solutions to this paradox invoke the emergence of karyotypic changes primarily in small populations, where genetic drift overrides natural selection (Bush et al., 1977), or that meiotic drive facilitates the spread of novel chromosomal variants (White, 1968). Alternatively, nearly neutral chromosomal rearrangements may accumulate in a stepwise manner, only becoming strongly underdominant in combination with each other (see Rieseberg, 2001). Since neither of these proposed solutions have received satisfactory empirical support, however, the importance of speciation by the classical model of karyotypic hybrid dysfunction remains disputed (Berdan et al., 2023; Potter et al., 2017).

Nevertheless, a strong correlation between rates of chromosomal evolution and speciation have been demonstrated in mammals (Bush et al., 1977), lizards (Leaché et al., 2016), and butterflies (De Vos et al., 2020). Furthermore, Augustijnen et al. (2024) modeled karyotype evolution in a phylogenetic framework, suggesting that chromosomal rearrangements predominantly coincided with, rather than emerging as a by-product of, speciation events. There is also direct evidence of reduced gene flow on fissioned or fused chromosomes between karyotypically distinct lineages of, e.g., mice (Franchini et al., 2010; Giménez et al., 2013), shrews (Basset et al., 2006), and butterflies (Mackintosh et al., 2023).

To better understand the mechanisms driving karyotype evolution, and its role in speciation, we focus on guenons (tribe Cercopithecini), an African primate group containing ∼35 species that radiated over the past ca. 12 million years (Jensen et al., 2023; Lo Bianco et al., 2017). During the course of their evolution, guenons underwent a remarkable karyotypic diversification resulting in diploid chromosome numbers ranging from 48 to 72 (Moulin et al., 2008). The guenon evolutionary history is characterized by extensive ancestral hybridization (Ayoola et al., 2020; Guschanski et al., 2013; Jensen et al., 2023; Svardal et al., 2017; van der Valk et al., 2020), with gene flow occurring also between species with different chromosome numbers (Jensen et al. 2023; Detwiler 2019). This puts into question the role and strength of chromosomal differences as reproductive barriers. Using long-read sequencing and chromosome conformation capture, we assembled the genomes of eight guenon species with different karyotypes and complemented them with additional three publicly available guenon genomes. This dataset allowed us to dissect the chromosome evolution of guenon genome architecture at a previously unattainable resolution, and to study the role of chromosomal fissions as barriers to gene flow between karyotypically divergent lineages.

## Methods

### Dataset, sample collection and sequencing

DNA samples of eight guenon species were collected from primary fibroblasts maintained in the collection of cryopreserved living tissues and cells of vertebrates (RBCell collection, Muséum National d’Histoire Naturelle, Paris, Table S1). These species belong to four of the six guenon genera, *Allenopithecus, Allochrocebus, Erythrocebus* and *Chlorocebus*. Representative genomes from the remaining two genera (*Chlorocebus* and *Miopithecus*) were downloaded from public archives (see Table S2), thus providing us with a dataset of 11 guenon species, representing all genera and spanning the karyotypic diversity present in guenons. High molecular weight DNA extraction was performed with the Monarch HMW DNA Extraction Kit for Cells & Blood (New England Biolabs) and sequenced on the Sequel II instrument to obtain PacBio HiFi reads. Hi-C processing was performed using Dovetail Omni-C preps, with libraries subsequently sequenced on the Illumina NovaSeq6000 S4 system (150bp paired end).

### Genome assembly and scaffolding

Remaining adapter content in the PacBio HiFi reads was trimmed using HiFiAdapterFilt (Sim et al., 2022), and residual Illumina adapters in the Hi-C reads were trimmed with fastp (Chen et al., 2018). Diploid draft assemblies were then generated with Hifiasm (Cheng et al., 2022), using the Hi-C data for haplotype phasing. Due to phasing errors on hemizygous sex chromosomes, we noticed that X/Y-linked contigs in some instances were split between haplotypes. To resolve this, we identified sex-linked contigs by aligning them to the rhesus macaque reference genome (*M. mulatta*, Mmul_10), and transferred all X and Y-linked contigs to haplotype 1 and 2, respectively, prior to scaffolding.

To scaffold the draft assemblies, we first used the one-line command provided by the Omni-C pipeline (version 2021/07/20, available on https://github.com/dovetail-genomics/Omni-C) to map the paired Hi-C reads to the respective draft assembly, record all valid ligation events, remove read pairs that may be formed by PCR duplicates, and generate the final bam-file. The final output of the sorted alignment file was used as input for YaHS v1.1 run with default settings (Zhou et al., 2023).

### Manual curation and quality assessment

Hi-C contact maps were generated using juicer v1.22.01 (Durand et al., 2016). These were then loaded into Juicebox v1.11.08 (Dudchenko et al., 2018) and used to manually correct obvious scaffolding errors, such as misplaced or misjoined contigs and false inversions (Figure S1). We also aligned each scaffolded assembly (both haplotypes) to the Mmul_10 reference using minimap2/2.28-r1209 with the asm20 preset (H. Li, 2018), and generated a synteny map for each scaffold with custom Python and R scripts. These synteny maps were then used as guidance in the manual curation process: Wherever the Hi-C maps were inconclusive, we oriented and joined contigs to maximize collinearity with *M. mulatta*. The contiguity and quality of the final assemblies were evaluated with assembly_stats (Trizna, 2020), and BUSCO completeness with compleasm (Huang & Li, 2023) using the primates_odb10 database.

### Identification of alpha satellite repeats

We used SOS-DNA to investigate the proportion of alpha satellite repeat (AS) DNA in our assemblies (https://github.com/ARChE-Team/paper-SOS-DNA). We ran SOSmonomers using default parameters and the provided example reference monomer sequence which includes the most abundant AS repeats in two previously studied Cercopithecini genomes (Cacheux et al., 2018, 2016). The AS proportion for each species was calculated as the ratio between the number of bases identified as AS in the dataset to the total number of bases in the same dataset. This was done separately for the PacBio reads, complete draft assemblies, and chromosome-level scaffolds (see Results).

### Genomic synteny, fissions and fusions

For the synteny analyses, we only considered chromosome-level scaffolds of haplotype 1, including all autosomes and the X chromosome. We used minimap2 (H. Li, 2018) to align each genome to the *M. mulatta* reference genome, using the asm20 option. Next, a custom Python script was used to generate a synteny map in a progressive manner: The genomes were ordered based on the guenon phylogeny (Jensen et al., 2024, 2023), and alignment coordinates were translated from *M. mulatta* to each specific target genome (see Figure 1). Next, we used the same alignments to call fissions and fusions in a more precise manner.

**Figure 1.**
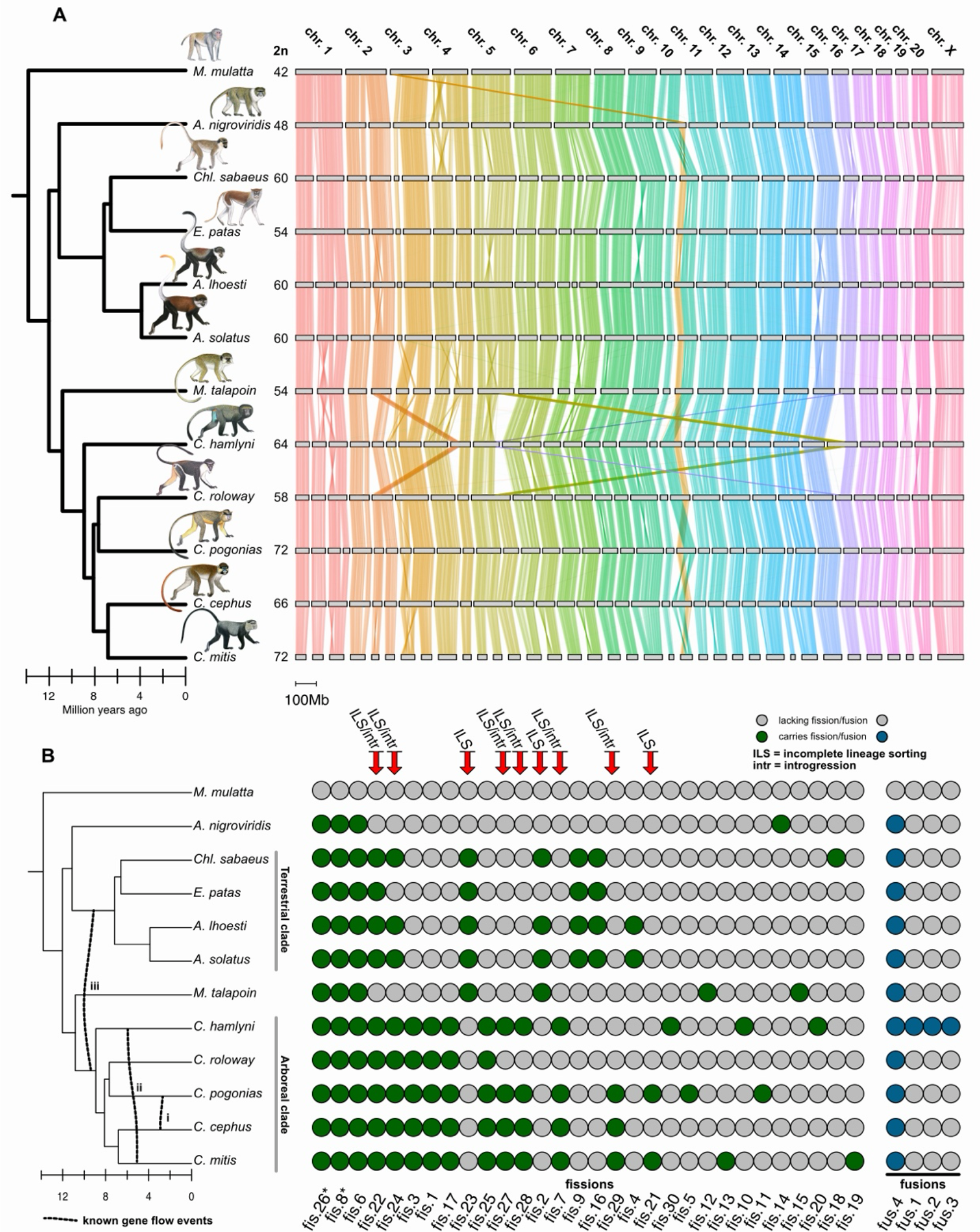
**(A)** Genomic synteny among guenons, with the species tree shown to the left (Jensen et al., 2024; 2023). Diploid chromosome numbers are displayed at the tips of the tree, with chromosomes shown as grey boxes and syntenic connections visualized as lines based on homology with the outgroup *M. mulatta* chromosomes. **(B)** Schematic overview of the 30 identified fissions and four identified fusions. The state for each fission/fusion (X-axis) in each species (Y-axis) is represented by gray (absent) or green/blue (present) circles. Species tree on the left is the same as in (A), with dashed vertical lines depicting known gene flow events (Jensen et al., 2023), see also main text. Red arrows highlight fissions that are discordant with the guenon phylogeny and thus result from incomplete lineage sorting (ILS) or introgression (intr). Arrow labels indicate indicate whether the discordance is consistent with introgression based on known gene flow events (ILS/intr) or more likely results from ILS alone (ILS). Primate illustrations copyright 2013 Stephen D. Nash/IUCN SSC Primate Specialist Group. Used with permission. *Fission 26 and 8 are present also in *H. sapiens*, suggesting that these likely result from fusion events in the outgroup (*M. mulatta*).

This was performed in two steps: First we used a custom Python script to identify sites in the genome assemblies where putative fissions or fusions occurred. A putative fission was defined as a shift in query scaffold but not reference chromosome along reference-sorted alignments. Fission breakpoint interval was identified as the highest reference coordinate in alignments left of the fission to the lowest reference coordinate in alignments on the right side of the fission. Putative fusions were called where the opposite pattern was found, i.e. a shift in reference chromosome but not query scaffolds. Since this approach may generate false positives due to misalignments or other, complex chromosomal rearrangements such as small translocations, the fissions and fusions for each species were manually curated using pairwise alignments as guidance. We then combined the genome-specific calls across all guenons, considering fissions and fusions with overlapping breakpoint intervals as single events. In complex events where the fission or fusion point was difficult to identify precisely, we used a parsimonious approach and manually combined events even if they did not show an exact overlap. For example, all guenons share a fusion event involving parts of the homologs to *M. mulatta* chromosome 3 and 10. However, the exact fusion point on chromosome 10 differs among species: In some it appears in the ancestral telomeric region, whereas in others the fusion occurred at a derived fission breakpoint closer to the center of the ancestral chromosome 10 (see Figure 1). We interpret this as a single event, but acknowledge the possibility of more complex scenarios with repeated fissions or subsequent translocations.

### Annotating repeats and segmental duplications

To characterize the repeat landscape of each genome, we used the EarlGrey pipeline (Baril et al., 2024). EarlGrey uses several open-source tools, including RepeatModeler and RepeatMasker, to generate *de novo* repeat annotations for genome assemblies. To explore the content of transposable elements (TEs) and repeats around chromosomal breakpoints, we first summarized the repeat content in 100 kb non-overlapping genomic windows for each genome using the R package GenomicRanges (Lawrence et al., 2013). For each fission event, we then translated the genomic coordinates of the breakpoint from the *M. mulatta* genome to the guenon species that retained the ancestral state (i.e. lacked the fission), using the liftover tool in paftools on the alignments to *M. mulatta*. We then visualized the proportion of the major repeat families along the outgroup homologs of the fissioned chromosome in proximity to the breakpoint (± 1 Mb).

We used biser/1.4 (Išerić et al. 2022) to characterize segmental duplications in the included genome assemblies. The genomes were softmasked prior to running biser, using the repeat annotations from EarlGray as well as windowmasker as implemented in the Galaxy toolshed (https://github.com/galaxyproject/tools-iuc/tree/main). From the raw calls, we removed candidates if there was an overlap in genomic position among mates with more than 20%, more than 70% of the base content was softmasked, or if the alignment error between mates exceeded 20%.

Finally, TRASH (Wlodzimierz, Hong, and Henderson 2023) was used to identify repeats specifically associated with centromeres. TRASH was run with default parameters, and consecutive regions were merged using bedtools (Quinlan and Hall 2010).

### Genome annotations and gene losses among guenons

To annotate the gene content of the genome assemblies and identify genes with loss of function mutations in guenon lineages, we used the pipeline TOGA (Kirilenko et al. 2023). TOGA is an orthology-based annotation tool, that identifies genes in query genomes based on a reference and classifies them as intact or lost based on the presence of inactivating mutations. We used the *M. mulatta* genome and annotation as reference, and a gene was considered lost in a species only if both haplotypes were annotated as ‘clearly lost’ by TOGA. For X-chromosomal genes, we only considered haplotype 1 (the “maternal” haplotype) in male samples. We excluded the publicly available highly fragmented *M. talapoin* and *C. mitis* genomes from this analysis.

### Gene flow

To test if chromosomal fissions and fusions acted as barriers to gene flow, we first used the alignments to the *M. mulatta* genome to call single nucleotide polymorphisms across all guenons. We used the ‘call’ function in paftools to call haploid genotypes for all assemblies against the *M. mulatta* reference, which were subsequently combined per species into diploid genotypes.

Next, we used ABBABABAwindows.py to estimate the *f*_*dM*_ statistic (https://github.com/simonhmartin/genomics_general; Malinsky et al., 2015; Martin et al., 2015) in 25 kb, non-overlapping windows along the *M. mulatta* chromosomes (Y-chromosome excluded). The *f*_*dM*_ statistic is a D-statistic derivative (Patterson et al., 2012), and quantifies excess allele sharing between two taxa based on a four-taxon topology. This test requires an outgroup, two monophyletic sister lineages (P1 and P2) and a midgroup (P3). A positive *f*_*dM*_ value indicates excess allele sharing (indicative of gene flow) between P3 and P2, whereas a negative value suggests excess allele sharing between P3 and P1. Based on Jensen et al. (2023), there are three known gene flow events among the species included in this study: between i) *C. cephus* and *C. pogonias*, ii) *C. hamlyni* and *C. mitis*, and iii) the *Cercopithecus* ancestor and the ancestor of the genera *Erythrocebus, Allochrocebus* and *Chlorocebus*, often referred to as the terrestrial clade (Tosi et al., 2004). We set up the analyses based on the guenon phylogeny and gene flow strength inferences from Jensen et al. (2023), using *C. roloway, C. cephus*, and *A. nigroviridis* as P1 lineages, respectively (see Figure 1B), and using *M. mulatta* as the outgroup.

We excluded windows with less than 100 SNPs, and used paftools liftover and the alignments to the *M. mulatta* reference genome to translate the genomic coordinates to the P2 species genome. This allowed us to assess *f*_*dM*_ on P2 chromosomes that were involved in fissions or fusions compared to those which retained a 1:1 orthology towards P3.

## Results

### Sequencing, genome assembly, scaffolding and manual curation

We generated PacBio HiFi and Hi-C sequencing data for the eight novel guenon genomes, achieving a median diploid depth of 37.4X (28.2 - 68.9) and 85.3X (68.9 -182.8), respectively (Table S1). Median PacBio HiFi read length was 16.7 kb (14.3 - 19.3). After scaffolding and manual curation of initial phased diploid assemblies, we obtained highly contiguous assemblies with a median contig N50 of 6.86-54.26 Mb (Table S2). The estimated sizes of the complete assemblies ranged from 2.83 to 3.84 Gb, with several being substantially larger than most available primate reference genomes, which are typically around 3 Gb (e.g. *Homo sapiens*: 3.1 Gb [GCF_000001405.40], *Macaca mulatta*: 3.0 Gb [GCF_003339765.1], *Chlorocebus sabaeus* 2.9 Gb [GCF_015252025.1]). The unexpectedly large assemblies contained many small, unplaced contigs (Figure S1), which were almost entirely made up of alpha satellite (AS) DNA (Figure S2). AS are the main components of centromeric DNA in Catarrhini, contributing up to 20% to the total genome size in some guenon species (Kurnit & Maio, 1973). The complete assemblies contained 0.37-19.17 % AS DNA (median 9.28 %). This is similar to the proportion of AS in the PacBio HiFi sequencing reads (Figure S3), which provide an unbiased estimate of the AS content, unaffected by the assembly process. In contrast, the proportion of AS across chromosome-level scaffolds in the assemblies was substantially lower for each species (0.32-4.03 %, median 2.28 %, Figure S3). This suggests that there indeed is a substantial genome size variation among guenons, primarily driven by centromeric DNA that we were unable to scaffold onto chromosomes. Since these unplaced scaffolds/contigs are difficult to align, and their true genomic positions are unknown, we excluded short unplaced contigs from downstream analyses.

All our assemblies contained the expected number of chromosome-level scaffolds, based on cytogenetically inferred karyotypes (Table S1; Moulin et al., 2008), and we thus refer to them as chromosome-level assemblies. By comparing our assemblies to cytogenetic images from Cacheux (2016) we were also able to map all chromosome-level scaffolds to cytogenetically identified chromosomes. BUSCO scores confirmed high completeness of the chromosome-level assemblies (> 98 % single Buscos; Table S2).

We complemented our eight newly generated genomes with the publicly available *Chlorocebus sabaeus* reference genome (GCF_000409795.2, Warren et al., 2015), and draft assemblies from *C. mitis* and *M. talapoin* (DNA Zoo Consortium; Dudchenko et al., 2017) (Table S2). We also included the *Macaca mulatta* reference genome (Mmul_10, GCA_003339765.3), to be used as outgroup and reference in our analyses.

### Guenon chromosomal evolution is dominated by fissions

To explore the chromosomal synteny among guenons in detail, we constructed a genome-wide synteny map and identified fission and fusion events. The genomic synteny map was constructed based on the guenon species tree (Figure 1A; Jensen et al., 2024, 2023).

Pairwise alignments to the *M. mulatta* genome were then used to identify shared and private fissions/fusions (Figure 1B).

In line with cytogenetic analyses (Dutrillaux et al., 1980; Moulin et al., 2008), we found that chromosomal fission was the main process behind the karyotypic differences in guenons. Across all included guenon genomes, we identified 30 putatively independent fissions and four fusions compared to the macaque (Figure 1B). Three fissions and one fusion, involving the homologs of *M. mulatta* chromosomes 2, 3 and 10, were shared among all guenon species. Two of these fissions (fission 8 and 26) were present also in the human genome (Figure S4), suggesting that they rather represent fusions occurring in the *M. mulatta* lineage. However, the other two events (fission 6 and fusion 4) were unique to guenons and thus likely occurred in their ancestor. *Cercopithecus hamlyni* was the only guenon species with additional species-specific fusions (Figure 1B). However, two of these three fusions likely result from a reciprocal translocation involving the homologs of *M. mulatta* chromosomes 5 and 16 (Figure 1A), and thus represent a single event.

Of the 21 *M. mulatta* chromosomes, only seven autosomes (chr. 9, 13, 15, and 17-20) and the X chromosome retained a 1:1 orthology with guenons. Despite the high rate of chromosomal fissions, intrachromosomal translocations and inversions were sparse.

We also note that the genomes of *M. talapoin* and *C. mitis*, which appear to harbor more inversions and translocations (Figure 1A), were assembled using short-read Illumina and Hi-C data (DNA Zoo consortium; Dudchenko et al., 2017), and are thus more prone to misassemblies compared to long-read-based assemblies used for the other guenon species. Despite the karyotypic diversity among guenons, genome collinearity is thus well preserved.

Nine of the identified fissions contradicted the phylogenetic tree, being paraphyletically distributed in the study species (Figure 1B). Such discordant patterns could arise from three alternative processes: independently reoccurring fissions (homoplasy), stochastic segregation of derived and ancestral chromosomal states among descending lineages (incomplete lineage sorting; ILS), or introgression of fissioned chromosomes through hybridization. Although studies propose that certain genetic features predispose chromosomal aberrations such as deletions, duplications and inversions (Durkin & Glover, 2007), which could possibly lead to repeated occurrences of such events, it is unclear if the same is true for fissions. Notably, non-centromeric chromosomal fissions, which appear most common in guenons (see below), also require the formation and activation of novel centromeres, making repeated events unlikely (Moulin et al., 2008).

Guenons experienced extensive ancestral hybridization in their evolutionary past (Jensen et al., 2023), and it is possible that fissioned chromosomes were transferred between species through introgression. Out of nine fissions that are discordant with the guenon phylogeny, six show a pattern that is consistent with introgression based on known ancestral hybridization events (Figure 1B). The remaining three discordant fissions cannot be explained by the known gene flow events and, assuming that repeated independent fissions are rare, likely result from ILS. For example, fissions 23 and 2 are present in *Miopithecus talapoin* and terrestrial clade species, but not in *Allenopithecus nigroviridis* or any *Cercopithecus* species (Figure 1B). If ILS caused these discordances, the fissions must have occurred in the guenon ancestor and remained segregating in the *Allenopithecus*/terrestrial clade ancestors and in the *Miopithecus/Cercopithecus* ancestor until after these respective lineages diverged, remaining polymorphic for at least ca. 1 million years. Similarly, under ILS, fission 21 would have emerged in the ancestor of *C. mitis, C. cephus, C. pogonias* and *C. roloway*, and remained segregating for ∼1.5 million years until reaching fixation independently in *C. mitis* and *C. pogonias*.

### No correlation between rate of karyotype evolution and genomic landscape of repeats and structural variation

Repetitive DNA is often invoked as a driver of karyotypic evolution. Hence, we characterized the repeat landscape of all genomes included in this study. The overall repeat content was highly similar among all guenons and *M. mulatta*, and there was no correlation between any repeat family and the number of chromosomes (Figure 2, invisible in my file; Spearman’s correlation tests: p ≥ 0.16, Table S3, S4). There was also no clear relationship between chromosome number and alpha satellite content in the PacBio reads, even though the two species with the highest number of chromosomes also showed the highest AS content (Figure S3B).

**Figure 2.**
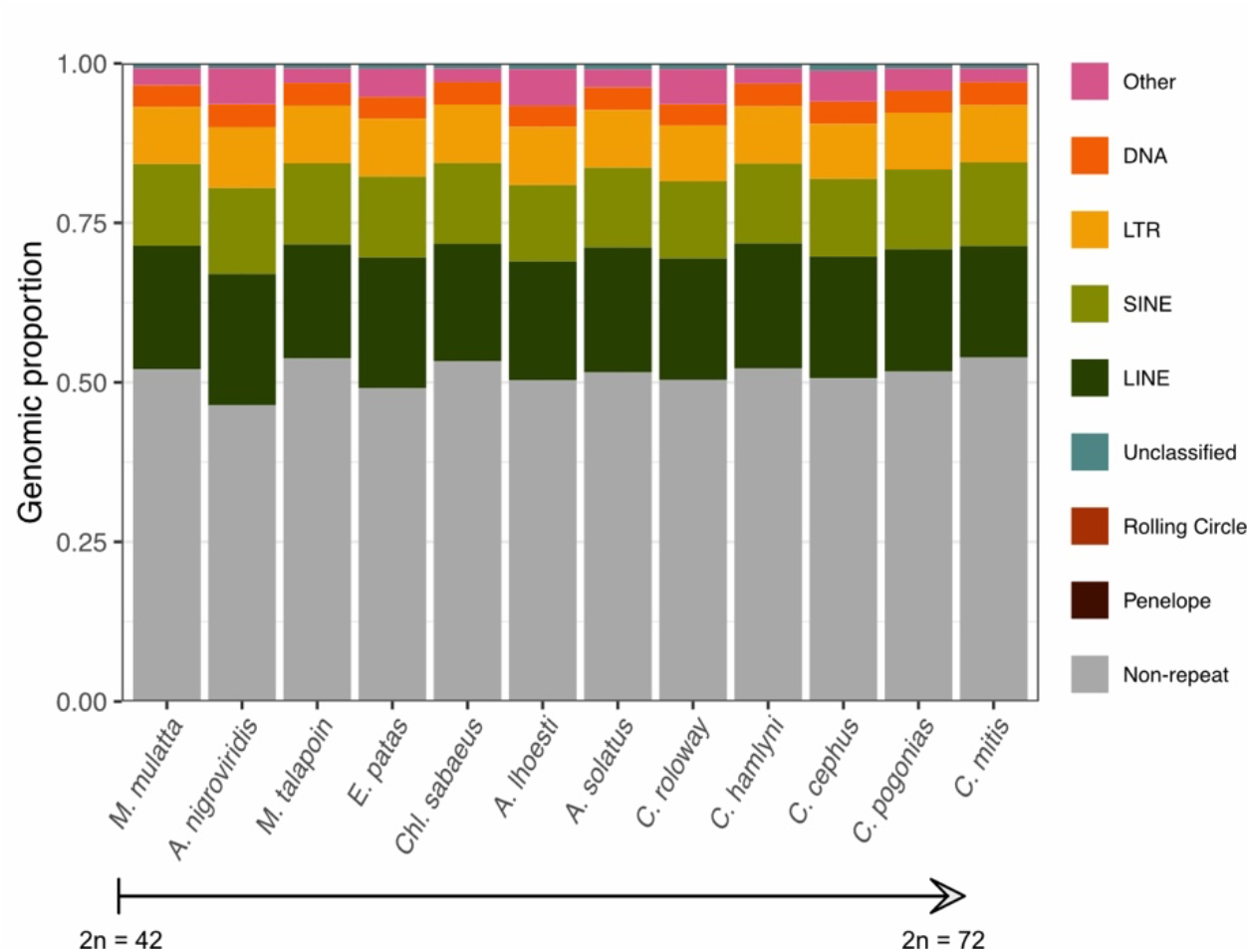
Repeat composition of guenon and *M. mulatta* genomes, arranged by ascending chromosome numbers from left to right.

If repeat expansions indeed were involved in the chromosomal instability of guenons, it must have been through fine scale processes, not detectable at the genome-wide level. Therefore, we also explored if fission breakpoints were enriched for specific repeat classes or segmental duplications, by characterizing the repeat content around the ancestral fission breakpoint in homologous, non-fissioned chromosomes (i.e., in guenon species lacking the specific fission, Figure S5-S30). Out of the 28 chromosomal fissions that occurred in guenons, two breakpoints (fission 14 and 18, private to *A. nigroviridis* and *Chl. sabaeus*, respectively; Figure 1B) could not be accurately inferred due to their proximity to complex inversions and/or translocations. Among the remaining 26 fissions, we identified five cases in which the breakpoints overlapped clear peaks of centromeric repeats in species without the fission, suggesting that these occurred within ancestral centromeric regions through centromeric (Robertsonian) breaks (Table S5; Figure S5-S9). One additional fission (fission 1, Figure S10) overlapped a putative centromere as inferred by TRASH in one of the outgroup species, again suggesting a centromeric fission. In contrast, 20 fissions were likely non-centromeric, with breakpoints occurring in regions away from putative ancestral centromeres (Figures S11-S30). Although three of these putatively non-centromeric breakpoints showed some evidence for expansions of segmental duplications and/or LINE repeat elements (fissions 3, 6 and 22, Figures S11, S14 and S25), the remaining 17 breakpoints did not show any association with repeat expansions. Overall, this suggests that repetitive elements were not direct drivers of the rapid rate of chromosomal fissions in guenons.

An alternative pathway by which repeat sequences and transposable elements (TE) could contribute to karyotypic diversity is by disrupting transcription of genes involved in chromosomal segregation, as was suggested to be the case in gibbons (Carbone et al., 2014). To explore this possibility, we used TOGA (Kirilenko et al. 2023) to classify genes that were disrupted by loss of function mutations in guenons compared to the *M. mulatta* reference genome. In total, TOGA identified 1,156 genes that were lost in any guenon lineage. Eleven gene ontology (GO) terms were enriched (FDR < 0.05) in this gene set, mostly related to sensory perception and smell (Table S6). In total, 180 genes were lost in all included guenon species, and 9 in all species except *A. nigroviridis* (i.e., lost in all species with accelerated karyotype evolution), but they showed no GO term enrichments.

Although several of the genes lost in any guenon species were annotated with GO terms related to chromosome organization or segregation, none of them were consistently lost in species with an accelerated rate of chromosomal fissions. Thus, our results do not support a scenario where disruption of chromosome segregation genes would be implicated in the guenon karyotype diversification.

### Reduced gene flow on fissioned chromosomes

Chromosomal rearrangements have been identified as barriers to gene flow in a number of species. We hence next explored whether fissions and fusions impeded ancestral gene flow, which occurred extensively throughout the guenon evolution (Jensen et al., 2023). To this end, we quantified local rates of introgression using the *f*_*dM*_ statistic (Malinsky et al., 2021; Martin et al., 2015) in 25 kb windows along the *M. mulatta* reference genome for three previously identified gene flow events (Figure 1B). The most pronounced event, with a gene flow proportion of ca. 15-20%, occurred between *C. cephus* and *C. pogonias* (event *i* in Figure 1B) (Jensen et al. 2023). Since diverging from *C. cephus, C. pogonias* experienced three additional fission events on the homologs of *M. mulatta* chromosomes 1, 3, and 5 (Figure 1, Figure 3A). We found that the *C. pogonias* autosomes deriving from these fission events showed reduced levels of gene flow compared to chromosomes that remained conserved between *C. cephus* and *C. pogonias* (Figure 3B; mean chromosome-wide *f*_*dM*_ 0.030 vs. 0.039, Wilcoxon rank sum test: p = 0.02). Since shorter chromosomes tend to show higher levels of gene flow (Edelman et al. 2019; Martin et al. 2019), likely due to higher recombination rate and hence reduced linked selection against introgression, we also ran a generalized linear model (GLM) with both synteny status (rearranged, conserved) and chromosome length as predictive variables. This test confirmed lower *f*_*dM*_ on fissioned chromosomes, independent of chromosome length (p_*synteny*_ = 0.01, p_*chromlength*_ = 0.13, Table S7). As expected, we detected a negative relationship, albeit marginally non-significant, between chromosome length and *f*_*dM*_ on conserved autosomes (Figure 3C; Pearson’s r = 0.36, p = 0.053). Our finding of a significantly reduced *f*_*dM*_ on rearranged chromosomes between *C. cephus* and *C. pogonias* suggests that fissions indeed act as barriers to gene flow.

**Figure 3.**
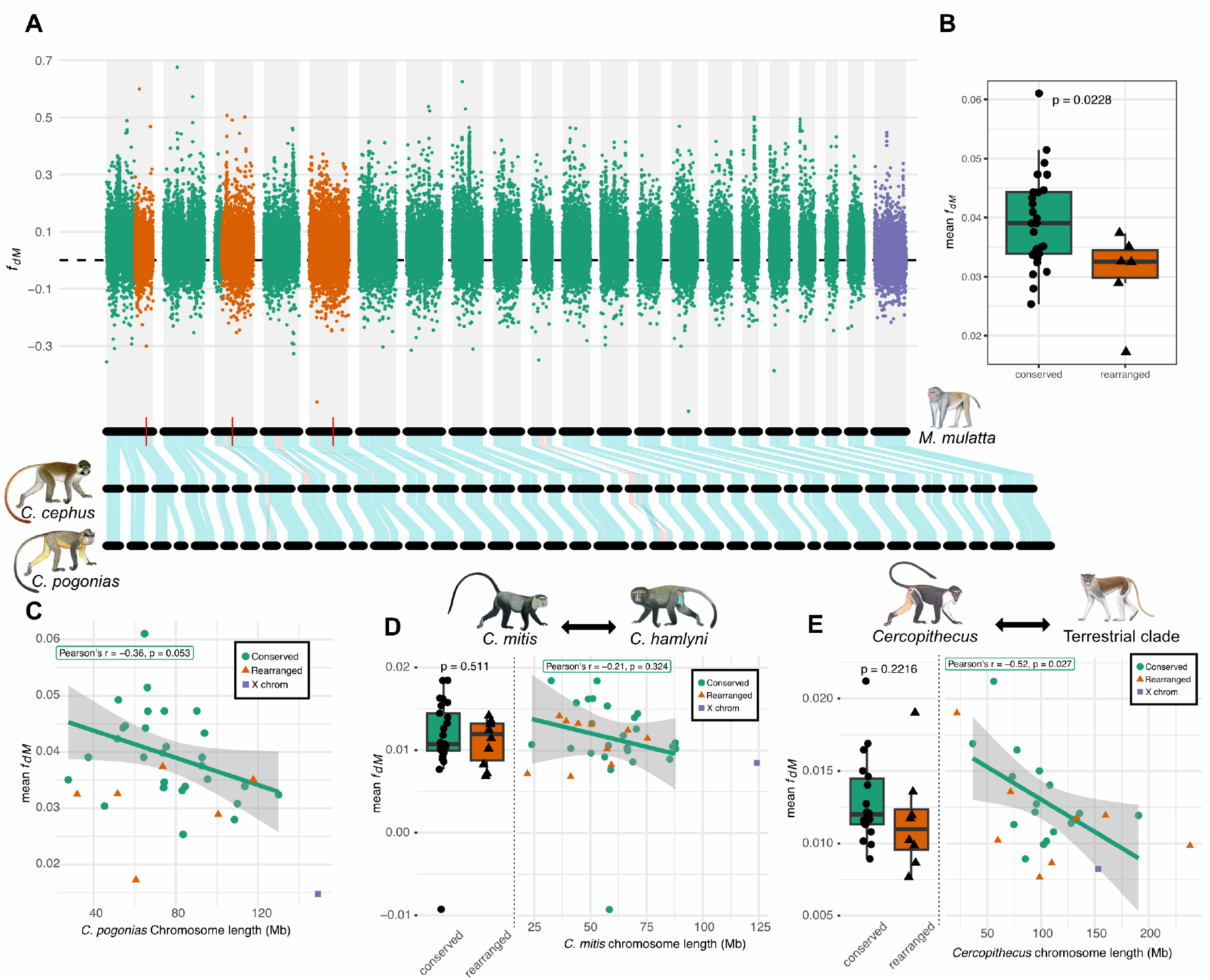
Genomic landscape of introgression. **A)** Local rates of introgression (*f*_*dM*_ estimates) in non-overlapping 25 kb windows along the *M. mulatta* reference genomes for the ancestral gene flow event between *C. cephus* and *C. pogonias*. Horizontal black bars below the plot illustrate the karyotypes of *M. mulatta, C. cephus* and *C. pogonias*, respectively. Vertical red lines on *M. mulatta* chromosomes highlight breakpoints of fissions private to *C. pogonias* relative to *C. cephus. f*_*dM*_ estimates on conserved autosomes between *C. cephus* and *C. pogonias* are shown in green, whereas orange points indicate a location on *C. pogonias* chromosomes involved in a private fission. X-chromosomal *f*_*dM*_ points are shown in purple. **B)** Average *f*_*dM*_ on conserved and rearranged *C. pogonias* autosomes, with p-value from Wilcoxon rank sum test. **C)** Average chromosomal *f*_*dM*_ (Y-axis) vs. *C. pogonias* chromosome length. Pearson’s correlation test was performed on conserved autosomes. **D)** Average *f*_*dM*_ between *C. hamlyni* and *C. mitis* on conserved and rearranged autosomes, including a p-value derived as in (B), and average *f*_*dM*_ plotted by chromosome length, with Pearson’s correlation test on conserved autosomes (right). **E)** Average *f*_*dM*_ between *C. hamlyni* and *C. mitis* on conserved and rearranged chromosomes, including a p-value derived as in (B), and average *f*_*dM*_ plotted by chromosome length, with Pearson’s correlation test on conserved autosomes (right).

Although we detected a tendency towards lower *f*_*dM*_ values on rearranged versus conserved chromosomes also in the two other known gene flow events that we were able to assess with included guenon taxa (events *ii* and *iii*, Figure 1B), this signal was not statistically significant (Figure 3D, 3E; p > 0.22; Table S7). These events also showed a negative relationship between chromosome length and mean *f*_*dM*_ on conserved autosomes, although it was only significant in the case of *Cercopithecus* and terrestrial clade ancestors (Figure 3E; Pearson’s r = -0.52, p = 0.027). In line with previous findings suggesting that sex chromosomes are often shielded from introgression (Fraïsse and Sachdeva 2021; Foley et al. 2026), the X chromosome showed among the lowest levels of *f*_*dM*_ of all chromosomes in all gene flow events.

## Discussion

We present high quality, chromosome-level genome assemblies from eight species of guenons with different karyotypes. Previous work on the guenon karyotype evolution was limited to cytogenetic studies (Dutrillaux et al., 1988; Milioto et al., 2023; Moulin et al., 2008). The genomes presented here thus offer the possibility to study the drivers of the remarkable chromosome evolution in guenons, and its role in speciation, at a previously unprecedented resolution.

When inferring the guenon phylogeny based on karyotypic traits, Moulin et al. (2008) retrieved a topology that differed substantially from the guenon species tree (Jensen et al., 2023). Indeed, we observed a high degree of discordance between guenon karyotypes and the species tree, as nine out of 28 fissions private to guenons did not follow the inferred evolutionary relationships. Chromosomal fission events are thought to be costly, as they require the formation and activation of a novel centromere and telomeres (Moulin et al., 2008). These discordances are thus arguably unlikely to be caused by recurrent independent fissions. Instead, incomplete lineage sorting (ILS) of polymorphic chromosomes likely generated this reticulate pattern, possibly in combination with introgression. Under ILS, fissioned chromosomes must have segregated in the ancestral guenon populations for extended periods of time, in some cases for more than one million years. In a similar scenario, a centromere position polymorphism on homologous chromosomes was proposed to have been segregating for several million years in ancestral guenon lineages, and retained polymorphic during multiple speciation events (Tolomeo et al., 2020). Tolomeo (2020) suggested that balancing selection was a likely driver of this long-lived polymorphism, which could also be the case for the pattern observed in our study. Regardless of any balancing selection, however, our results strongly suggest a weak or absent underdominance, i.e., negative selection against chromosomal heterozygotes. Alleles that cause underdominance are unlikely subjects of ILS, as they would be rapidly purged by selection (or possibly drift to fixation in small populations). Similarly, disadvantageous alleles are unlikely to be transferred between lineages through introgression, as they would be removed by selection in the recipient population.

The classical model of chromosomal speciation suffers from an important problem, as chromosomal variants that cause reproductive isolation between populations are unlikely to spread within the population due to underdominance. Several solutions to this paradox have been proposed, for example that chromosomal rearrangements predominantly arise in small populations, where genetic drift overrides selection. Following this logic, Bush et al. (1977) suggested that the social structure of guenons, which typically live in single-male/multiple-female groups, would lead to reduced effective population sizes making them more prone to chromosomal rearrangements than, e.g., karyotypically stable baboons (genus *Papio*), which form larger multi-male/multi-female groups. However, strong genetic drift should rapidly drive novel rearrangements to fixation, making ILS unlikely. Furthermore, recent demographic inferences show that guenons are among the most genetically diverse primates in the world, with large ancestral effective population sizes (Kuderna et al., 2023), which would imply that selection against deleterious variants should be efficient. Our results are thus not compatible with a scenario in which small effective population sizes were important drivers of the karyotype diversification in guenons. Instead, our findings suggest that fission events have been well tolerated in ancestral guenon populations, questioning their causal role in species diversification.

Nevertheless, we do find evidence of reduced gene flow on rearranged chromosomes in the most pronounced ancestral introgression event in guenons (between *Cercopithecus cephus* and *C. pogonias*, (Jensen et al. 2023)). A similar pattern, albeit not statistically significant, was also found in the gene flow event between the arboreal genus *Cercopithecus* and the terrestrial clade ancestors. Fissions and fusions were found to act as barriers to gene flow in several species (e.g. Basset et al., 2006; Franchini et al., 2010; Giménez et al., 2013; Mackintosh et al., 2023), suggesting that the model of chromosomal speciation through hybrid dysfunction may be important in some systems. Although the extensive phylogenetic discordance of chromosomal fissions in guenons suggests that single fission events have been well tolerated, it is possible that accumulation of several fissions generated barriers to gene flow (see Rieseberg, 2001), e.g., by increasing the rate of aneuploid gametes. Importantly, while our results suggest that chromosomal fissions likely acted as local barriers during post-speciation gene flow, their importance in the speciation process remains unclear. For instance, *C. cephus* and *C. pogonias* likely diverged ca. 3-4 million years prior to the hybridization (Jensen et al., 2024, 2023). Similarly, the genus *Cercopithecus* and terrestrial clade ancestors diverged more than one million years prior to the major gene flow event. Hence, reproductive isolation between these lineages may already have been strong, and the contribution of fissions to early stages of divergence is difficult to discern. Another important limitation is the uncertainty about the order of events, specifically if fissions pre- or post-dated gene flow. Fissions could only act as barriers to gene flow if they occurred prior to hybridization. We thus expect no signal of reduced gene flow on chromosomes that became rearranged after the occurrence of gene flow, which might explain the lack of significantly reduced introgression on rearranged chromosomes between *C. mitis* and *C. hamlyni*. Without knowledge about the karyotypes at the time of gene flow, we risk treating conserved chromosomes as rearranged, thus reducing our power to detect barrier effects. Therefore, we may have underestimated the reduction in gene flow on rearranged chromosomes in the three ancestral hybridization cases. Studying gene flow across contemporary hybrid zones between lineages with different karyotypes offers a solution to this problem. One such guenon hybridization zone is known from Tanzania, where *C. ascanius* (2n = 66) and *C. mitis* (2n = 72) overlap and hybridize extensively (Detwiler 2019), providing an excellent system for future studies on cross-karyotypic gene flow.

Why some lineages appear more prone to undergo chromosomal rearrangements, while others remain stable, is poorly understood. However, transposable elements (TE) have been implicated in several cases. For example, the chromosomal stability among birds has been attributed to their low levels of TE content relative to the more karyotypically diverse mammals (Ellegren, 2010). Furthermore, Carbone et al. (2014) demonstrated that lineage-specific TE insertions disrupted the function of important chromosome segregation genes in gibbons (family Hylobatidae), another primate lineage that experienced extensive chromosomal fissions. Our analyses did not reveal any obvious changes in repeat composition that could explain the chromosomal instability in guenons. Neither did we find any enrichment of chromosomal segregation functions among genes affected by loss of function mutations. The predisposition for guenons to undergo chromosomal fissions therefore remains a mystery at this stage, and elucidating the processes behind this extensive karyotypic diversification will require more research, combining cytogenetic and molecular methods.

In summary, this study provides a first glance at potential mechanisms of the rapid karyotype evolution in guenons and suggests that chromosomal fissions may have acted as barriers to gene flow during this primate radiation. It provides a foundation for future research into the mechanisms of chromosomal instability, origination of novel centromeres and telomeres, and the interplay between chromosome evolution and speciation.

## Acknowledgements

The computations and data handling were enabled by resources in projects NAISS 2023/6-340 and NAISS 2023/5-506, provided by the Swedish National Infrastructure for Computing (SNIC) at Uppsala University (UPPMAX), funded by the Swedish Research Council through grant agreement no. 2022-06725. The project was supported by the Swedish Research Council VR grant (2020-03398) to KG.

## Author contributions

Conceptualization: A.J. and K.G.; Methodology and analyses: A.J. with assistance from A.S., F.C., M.S., L.P., C.E., and J.P.; Sample acquisition: B.B., M.G-S., C.E.; Acquisition of funding: K.G.; Writing: A.J. and K.G. with input from all authors.

## Code availability

Scripts used in this project are available at https://github.com/axeljen/guenomics_2026.

## Data availability

Genome assemblies and sequencing data will be uploaded to public archives upon publication.

