## Supplementary figures for "Karyotypic evolution in a diverse primate clade and the role of chromosomal fissions as barriers to gene flow"

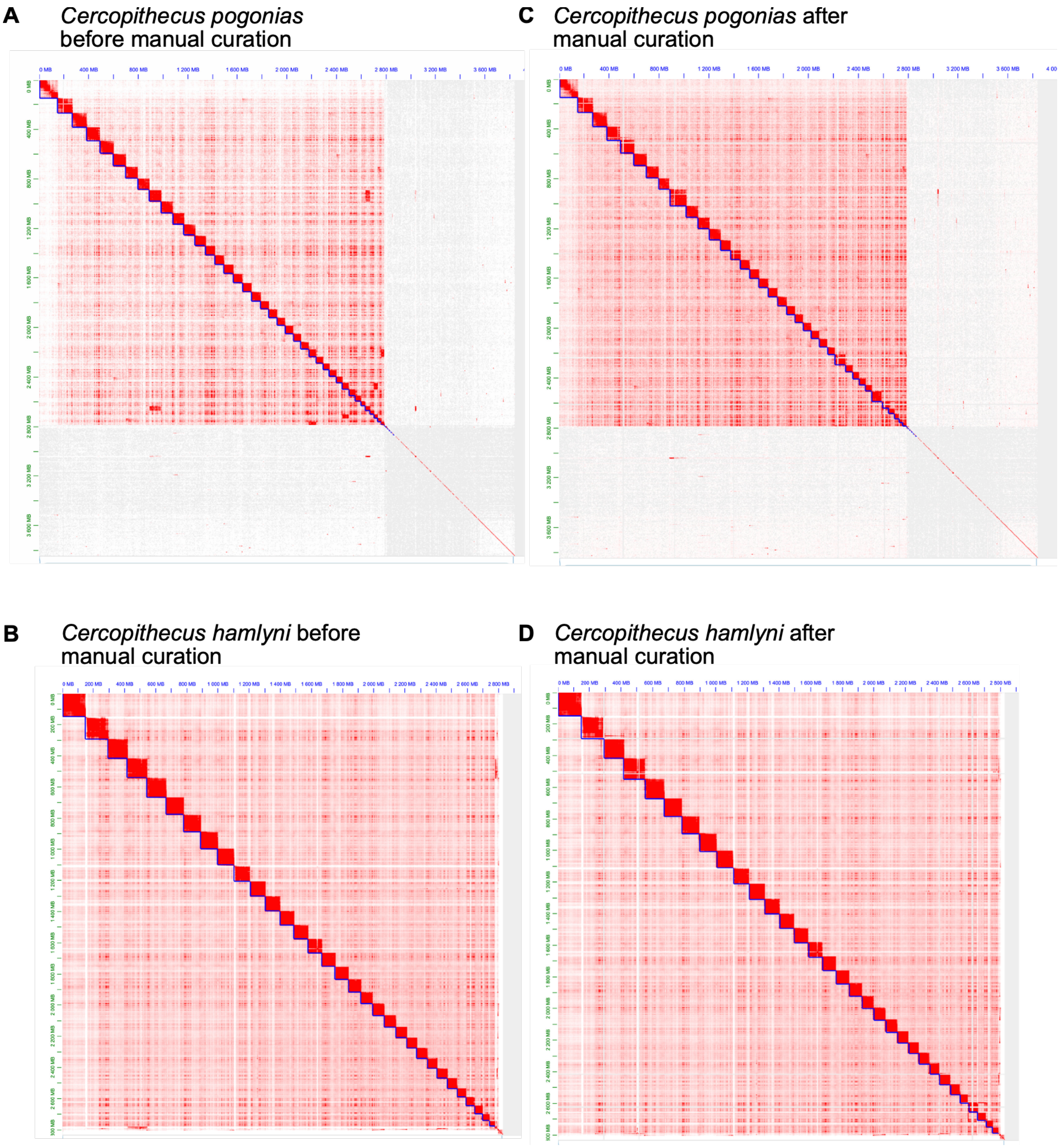


**Figure S1.** Examples of Hi-C heatmaps before (A,B) and after (C,D) manual curation for two species. Darker red indicates a high degree of chromatin interaction, as is expected for intrachromosomal, physically adjacent contigs. *Cercopithecus pogonias* had the largest draft genome size (3.84 Gb), attributed to many small contigs with poor Hi-C read mapping and thus low degree of chromatin interactions (lower right corner in A and C). *Cercopithecus hamlyni* had the smallest draft genome size (2.83 Gb), with the vast majority of contigs being placed on chromosome-level scaffolds (B, D).

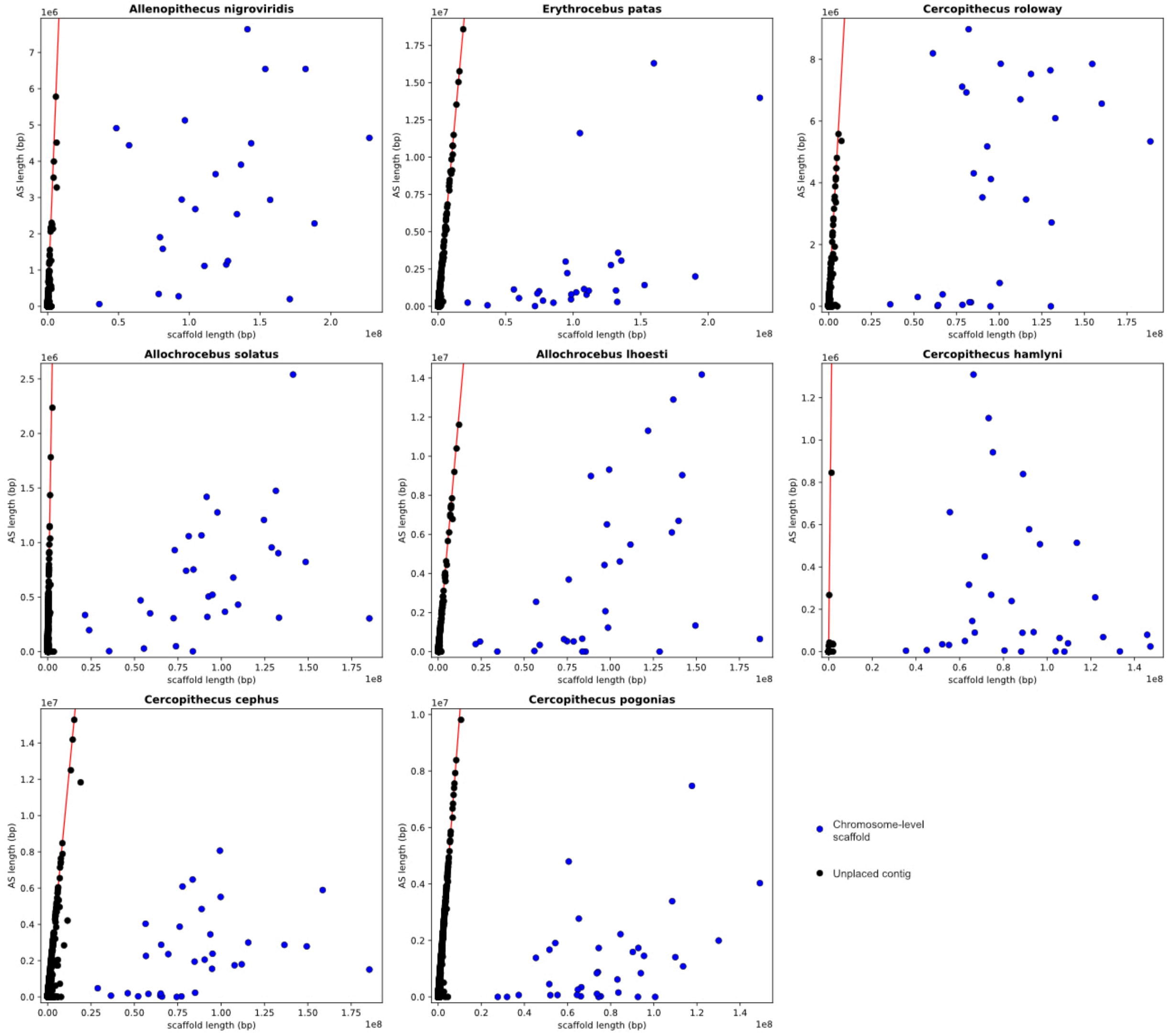


**Figure S2.** Summed length of alpha satellites (bp, *Y*-axes) in scaffold/contig sorted by length (bp, *X*-axes). Blue circles represent chromosome-level scaffolds (i.e., those that were retained in the chromosome-level assemblies), whereas black points show unplaced contigs. The 1:1 line is depicted in red.

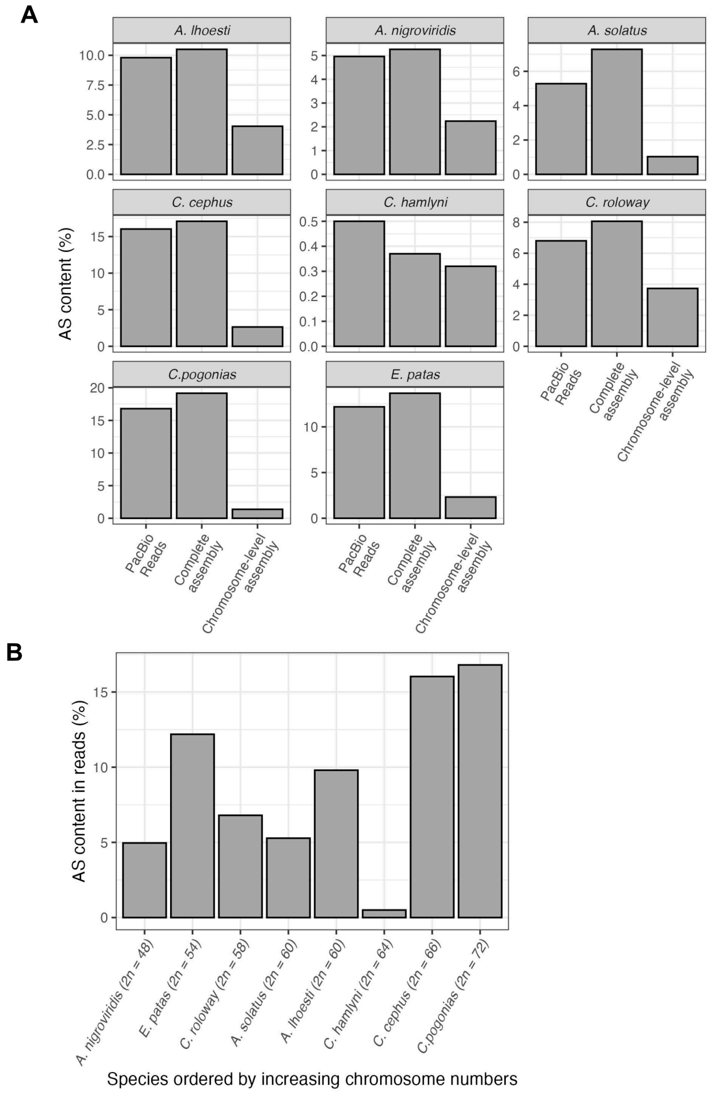


**Figure S3. (A)** Percentage of alpha satellite (AS) repeat content detected in the PacBio reads, total assembly (including unplaced contigs), and chromosome-level assemblies, for each species. **(B)** Percentage of AS content detected in PacBio reads per species, arranged by increasing chromosome numbers.

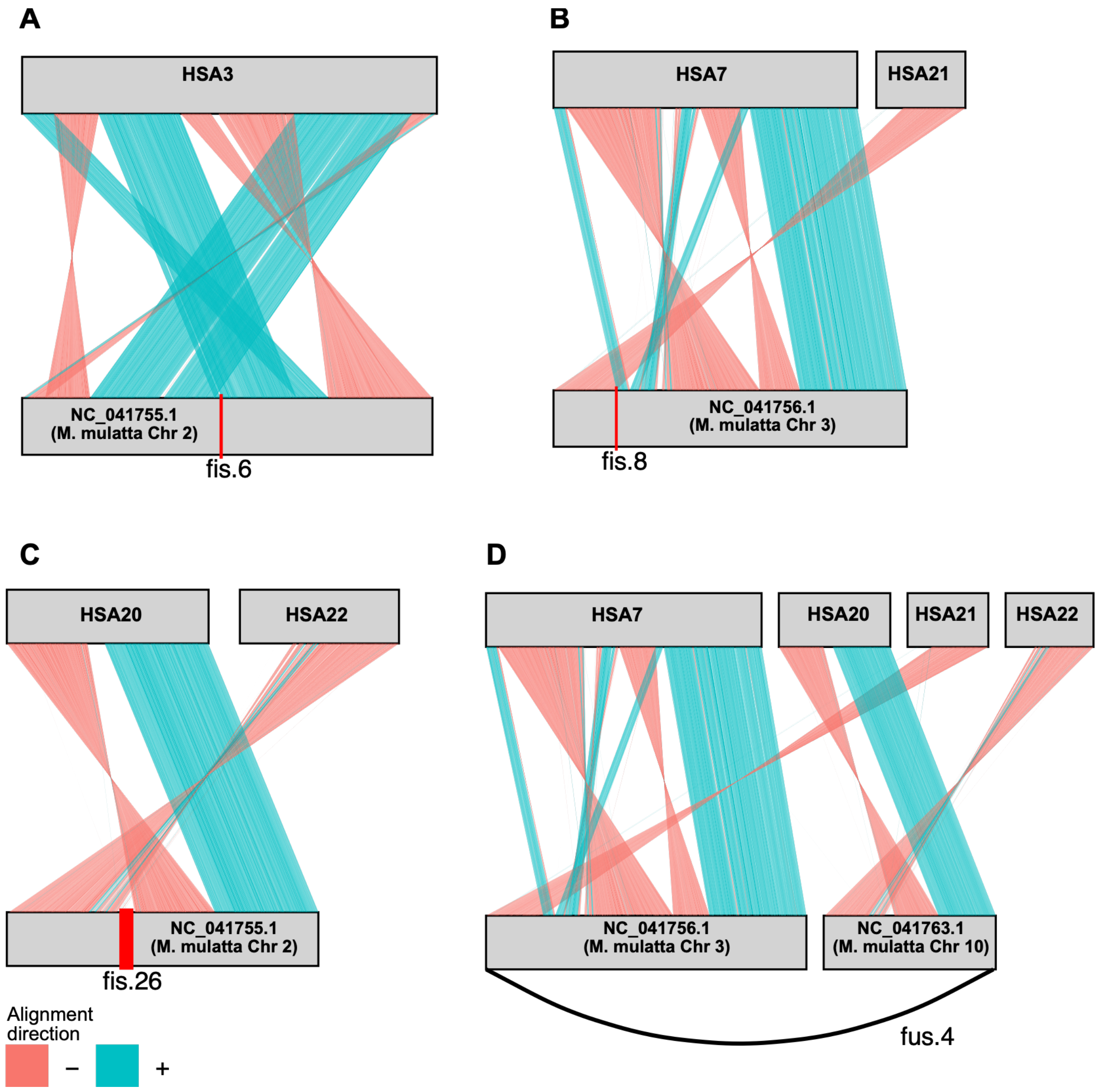


**Figure S4.** Homology between *M. mulatta* and *Homo sapiens* in the gproximity of three fission (A-C) and one fusion (D) that were shared among all guenons. The breakpoint of fission 8 and 26 (B and C, respectively) mark a chromosomal break also in the human genome, suggesting that these rather represent fusions occurring along the *M. mulatta* lineage. Fission 6 and fusion 4 appear to be private to guenons, suggesting an occurrence in their common ancestor.

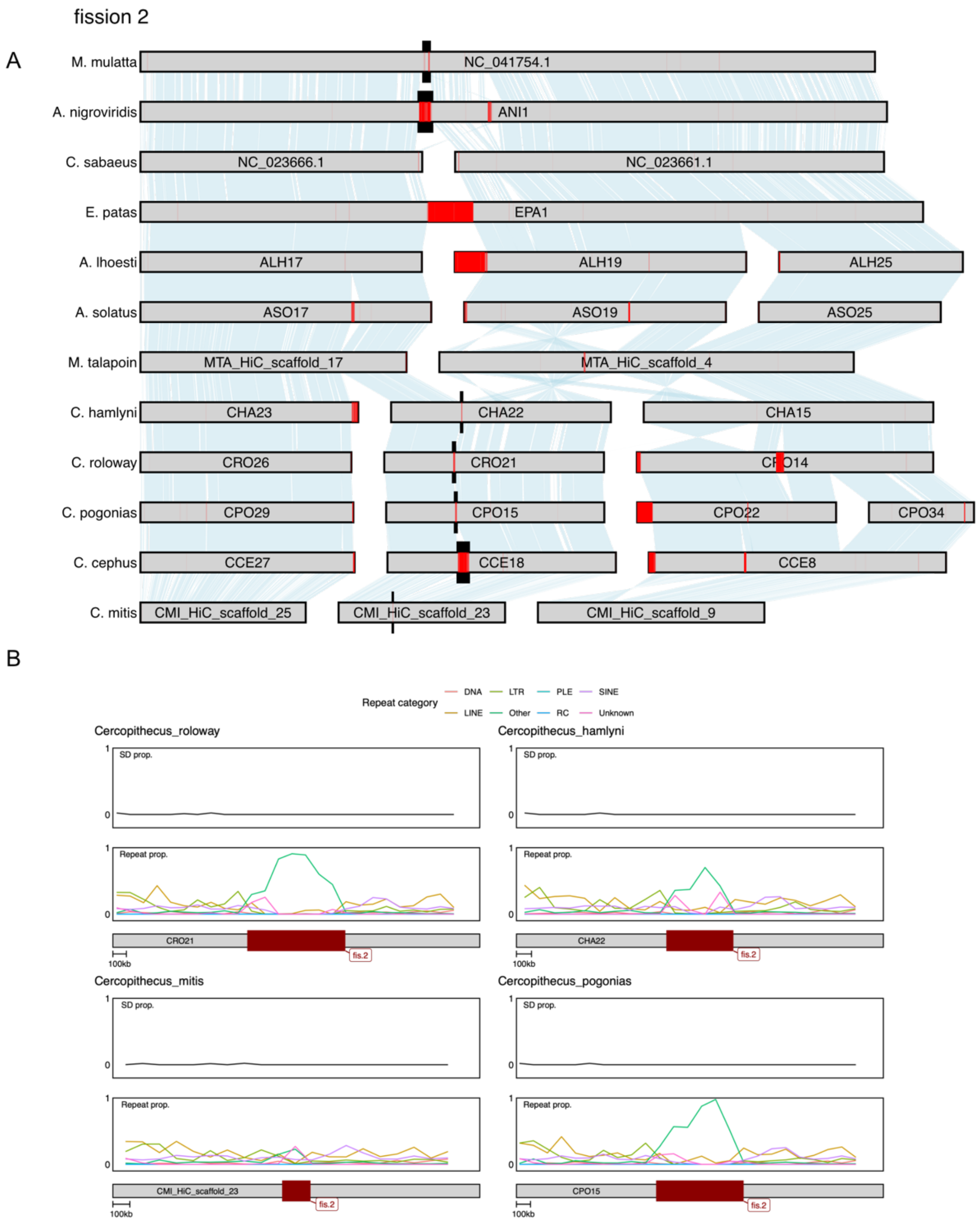


**Figure S5.** A) Chromosomal synteny around the ancestral breaking point of fission 2. Black markings highlights the fission breakpoint, and centromeric annotations from TRASH is highlighted in red. B) Proportion of segmental duplications (SD, top panels) and repetitive elements as annotated by EarlGrey (bottom panels) summarized in 100 kb sliding windows, in lineages lacking the fission. Fission 2 overlaps a putative ancestral centromere in the majority of species lacking the fission (A), and show clear peaks of repeat category "Other", which contains centromeric satellite repeats, from EarlGray in several outgroups (B). This fission thus likely represents a centromeric fission.

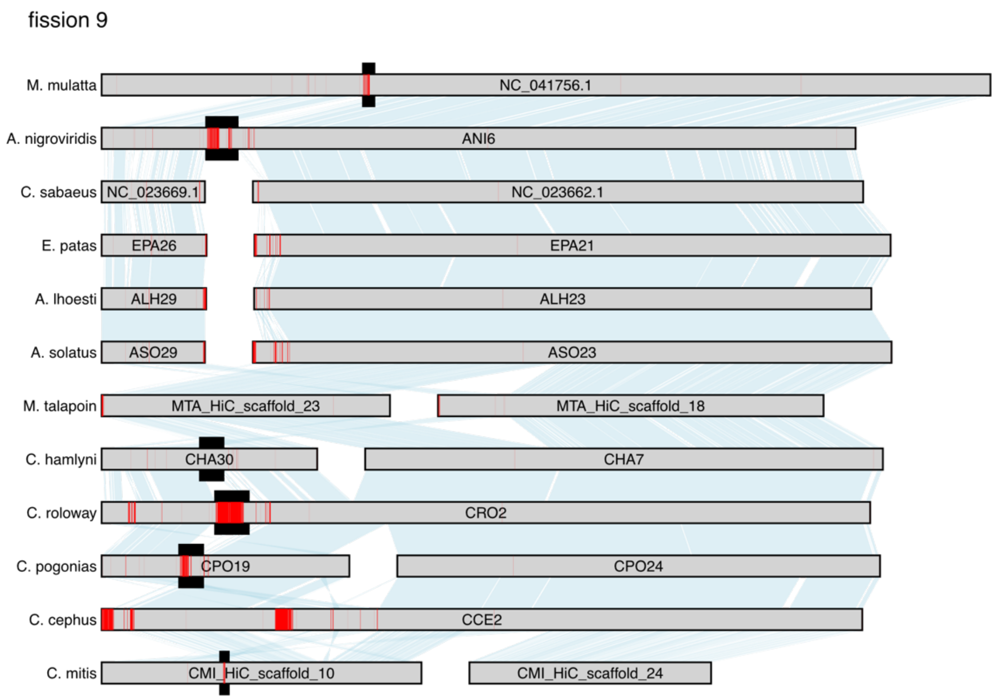


**Figure S6.** Chromosomal synteny around the ancestral breaking point of fission 9. Black markings highlight the fission breakpoint, and centromeric annotations from TRASH is highlighted in red. The fission overlaps putative centromeres in the majority of outgroups, suggesting that this event constitutes a centromeric fission. Breakpoint coordinates could not be inferred with sufficient accuracy for an analysis of SD and repeat content.

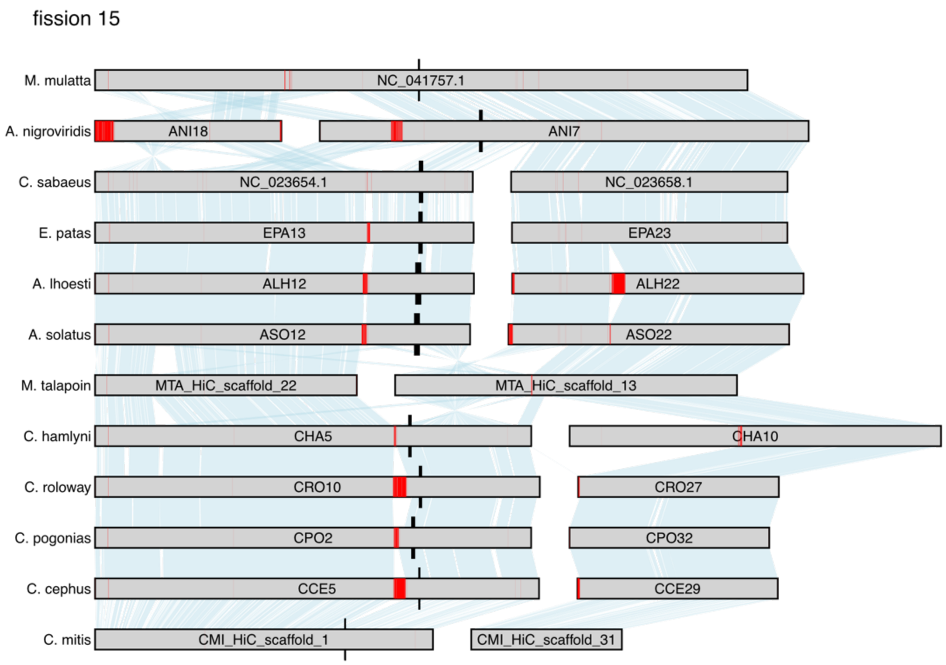


**Figure S7.** A) Chromosomal synteny around the ancestral breaking point of fission 15. Black markings highlight the fission breakpoint, and centromeric annotations from TRASH is highlighted in red. The fission beakpoints could not be accurately inferred due to complex rearrangements surrounding the breakpoints in *M. talapoin*, and the highlighted ancestral breakpoints are likely incorrect. However, the distal end of one of the fission products (MTA_HiC_scaffold_22) is homologous to a region containing a putative centromere in several outgroup species, and it is likely that the breakage occurred there, rather than at the highlighted regions in this figure (which is obscured by inversions and translocations).

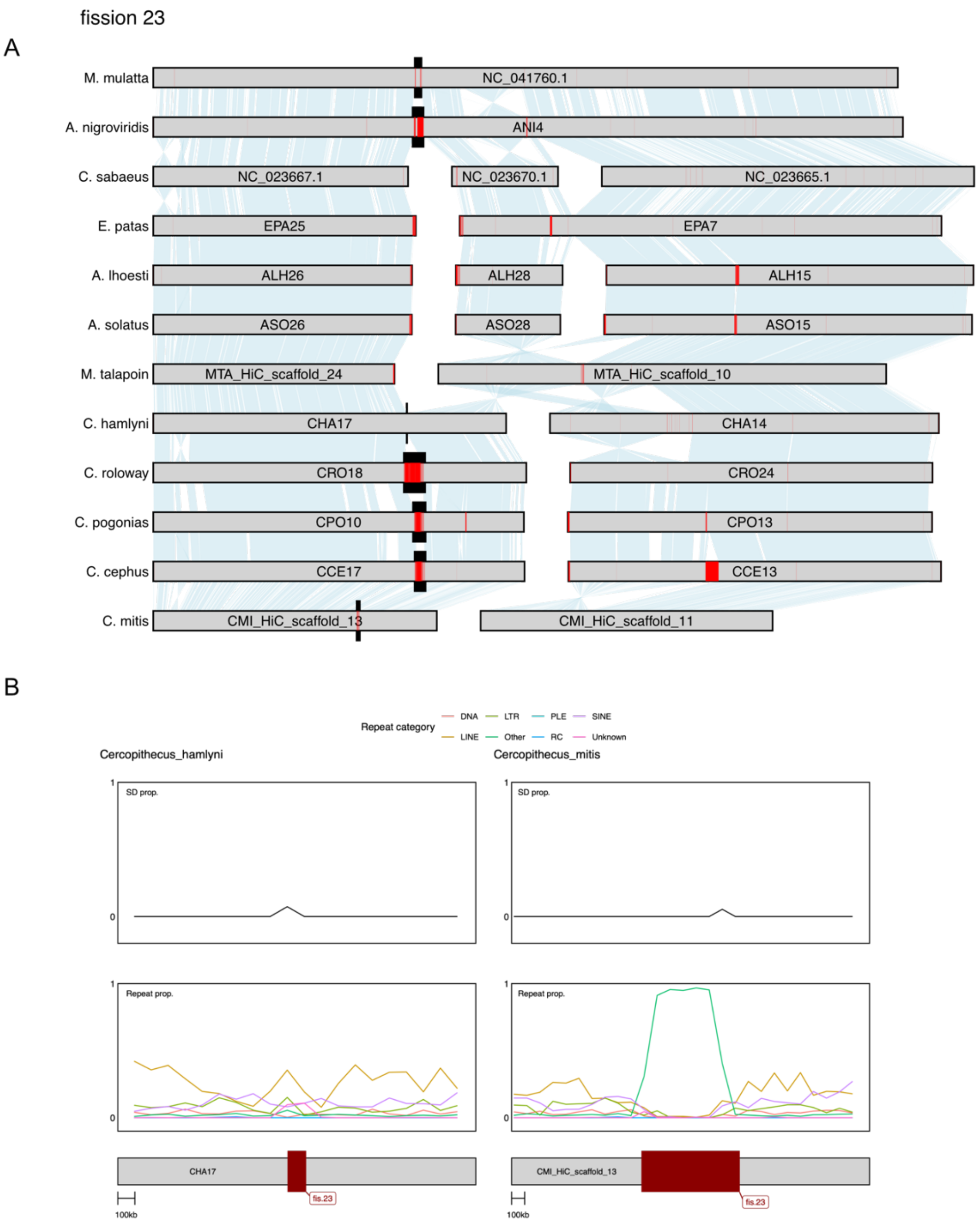


**Figure S8.** A) Chromosomal synteny around the ancestral breaking point of fission 23. Black markings highlight the fission breakpoint, and centromeric annotations from TRASH is highlighted in red. B) Proportion of segmental duplications (SD, top panels) and repetitive elements as annotated by EarlGrey (bottom panels) summarized in 100 kb sliding windows, in lineages lacking the fission. The fission overlaps putative centromeres in the majority of outgroup species, and a large peak of repeats in the category “Other”, which includes centromeric satellite repeats.

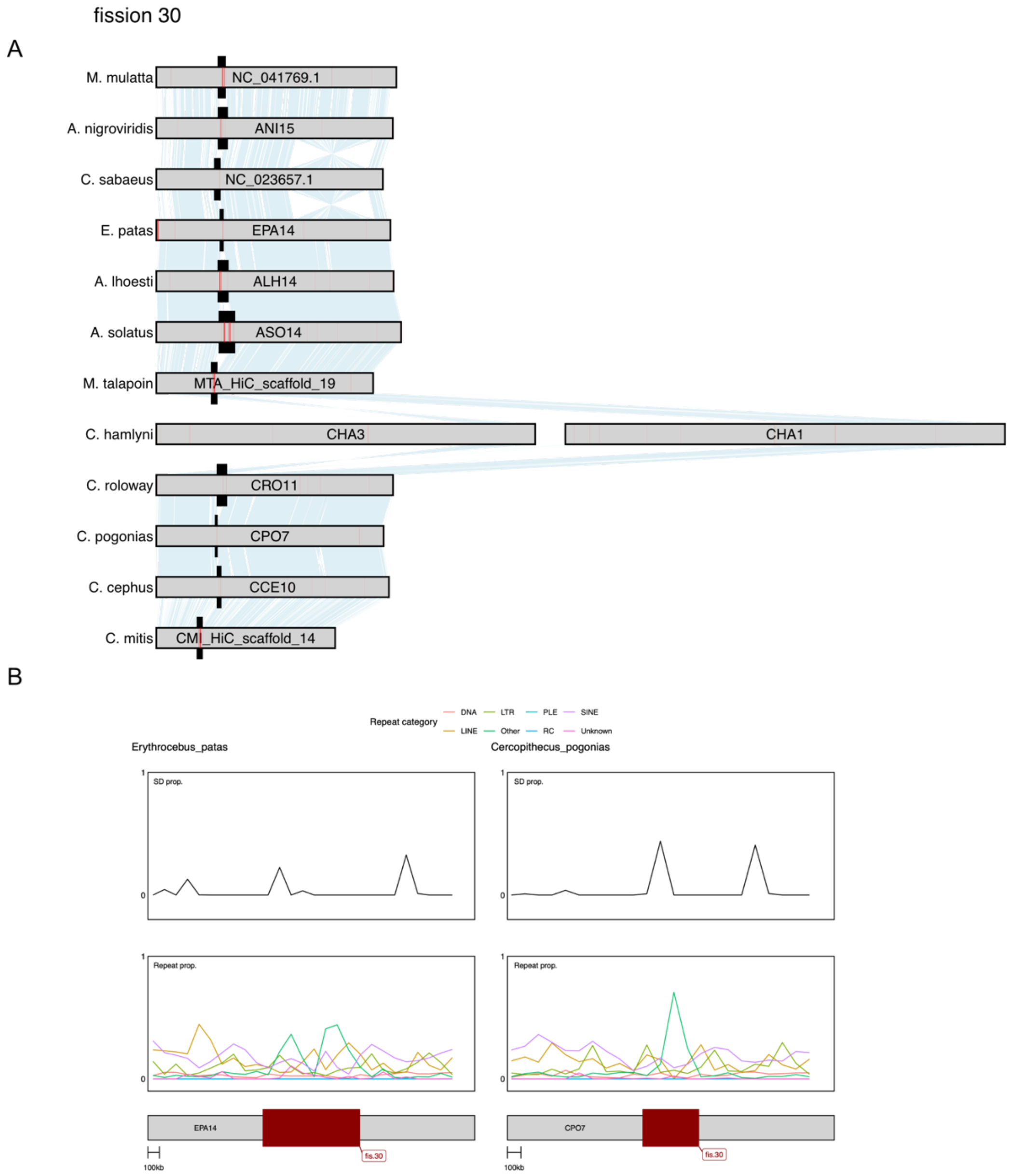


**Figure S9.** A) Chromosomal synteny around the ancestral breaking point of fission 30. Black markings highlight the fission breakpoint, and centromeric annotations from TRASH is highlighted in red. B) Proportion of segmental duplications (SD, top panels) and repetitive elements as annotated by EarlGrey (bottom panels) summarized in 100 kb sliding windows, in lineages lacking the fission. The fission breakpoints overlap any putative centromeres in the majority of outgroups. Breakpoint coordinates were too wide for meaningful inferences of local SD and repetitive element content in most outgroups, but some peaks of SD and "Other" repeat elements, including centromeric satellite repeats, were visible in the two outgroups where we could infer this (*E. patas* and *C. pogonias*)

fission 1

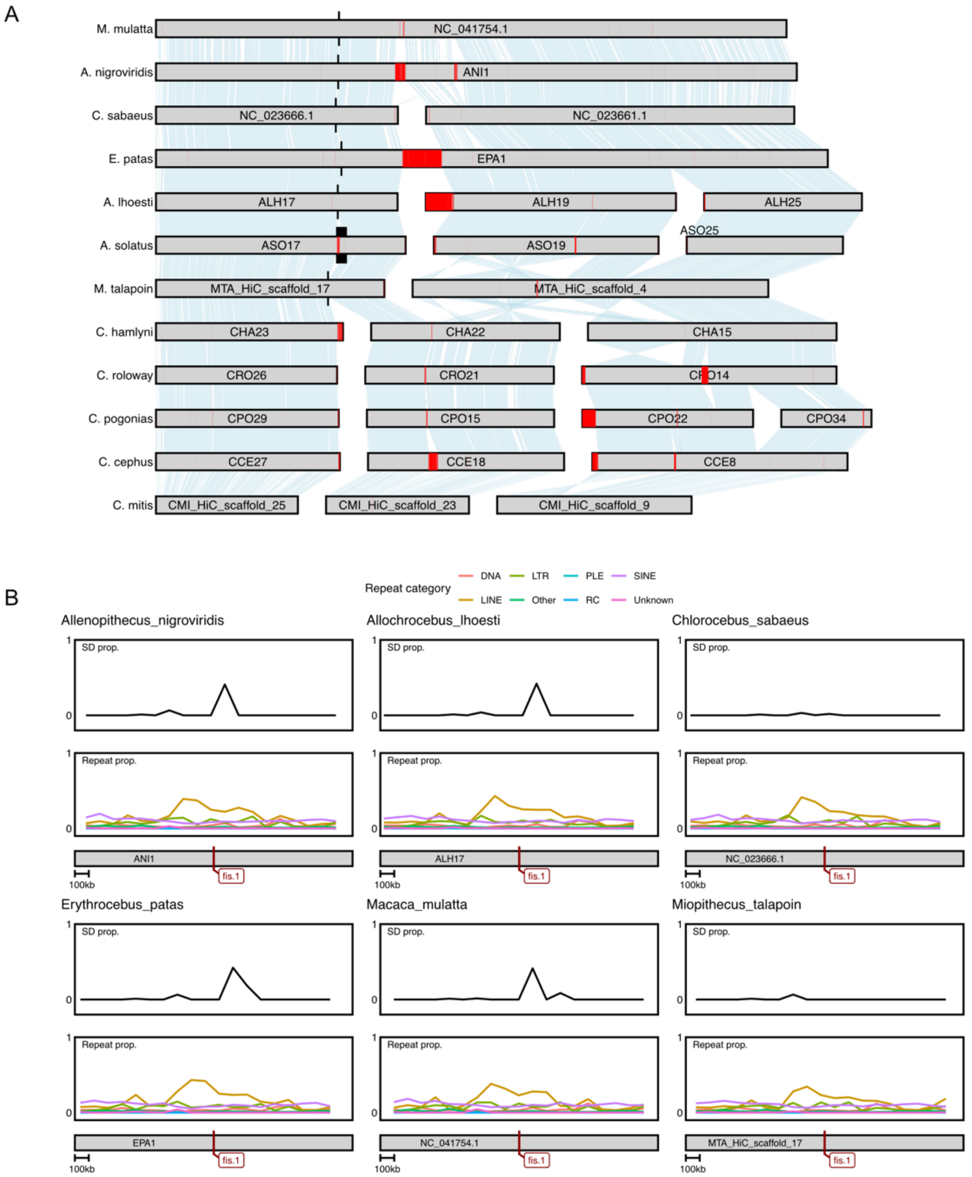


**Figure S10.** A) Chromosomal synteny around the ancestral breaking point of fission 1. Black markings highlights the fission breakpoint, and centromeric annotations from TRASH is highlighted in red. B) Proportion of segmental duplications (SD, top panels) and repetitive elements as annotated by EarlGrey (bottom panels) summarized in 100 kb sliding windows, in lineages lacking the fission. Fission 1 occurred far upstream the putatively ancestral centromere (A). Nevertheless, the breakpoint overlaps a region containing annotated centromeric repeats in one outgroup (*A. solatus*), and one of the fission products is likely acrocentric, with the centromere close to the breakpoint (*C. hamlyni* chr. CHA23). Furthermore, there is a slight increase of repetitive elements and SD content in several outgroups (species that do not carry the fission) around the breakpoint, which could be a hallmark of a centromere. It is therefore possible that this breakpoint occurred in a (derived) centromeric region.

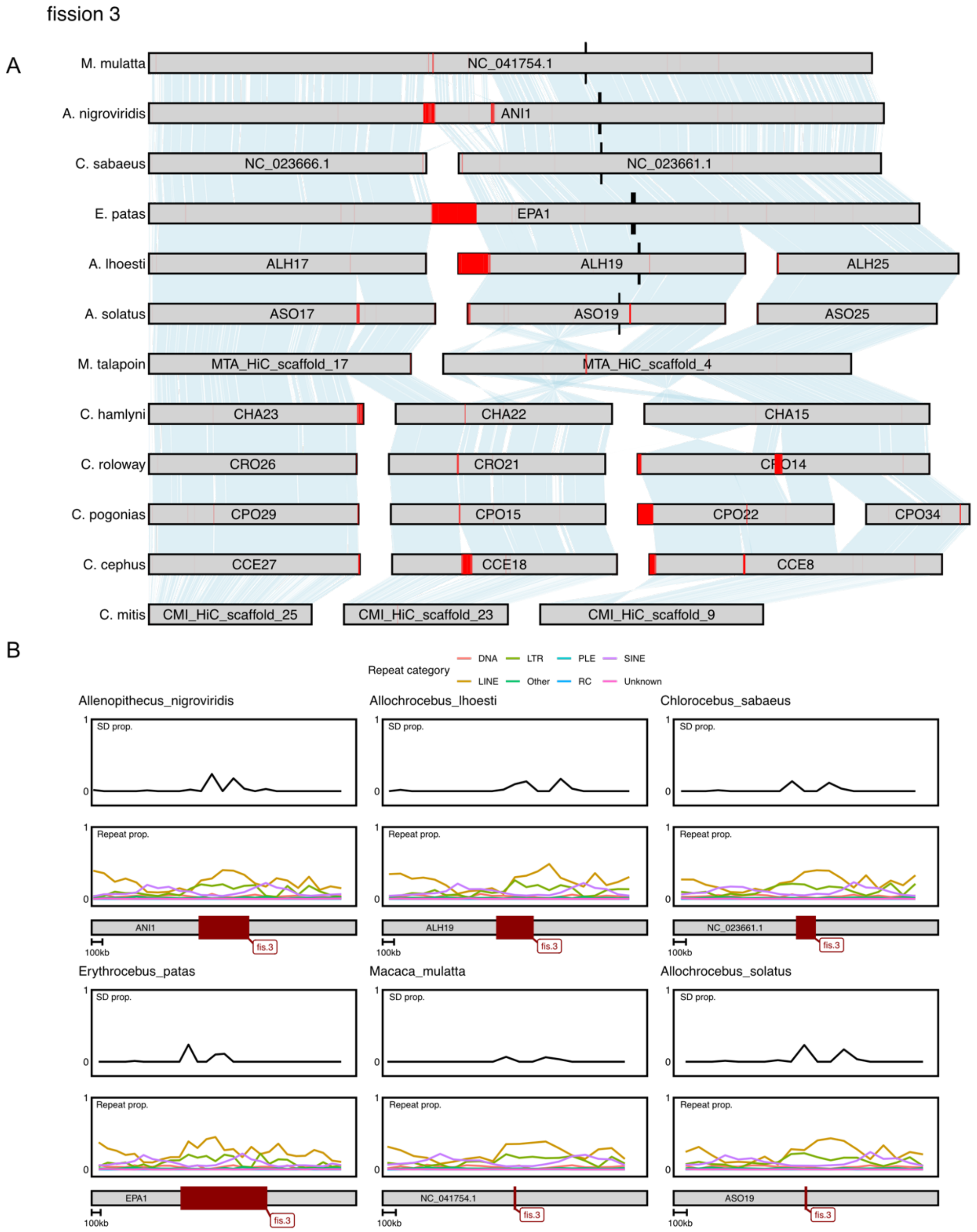


**Figure S11.** A) Chromosomal synteny around the ancestral breaking point of fission 3. Black markings highlights the fission breakpoint, and centromeric annotations from TRASH is highlighted in red. B) Proportion of segmental duplications (SD, top panels) and repetitive elements as annotated by EarlGrey (bottom panels) summarized in 100 kb sliding windows, in lineages lacking the fission. Fission 3 does not overlap putative centromeres in any lineage (A), suggesting this is a non-centromeric fission. Shallow peaks of SD content is seen around the breaking point all outgroups (B).

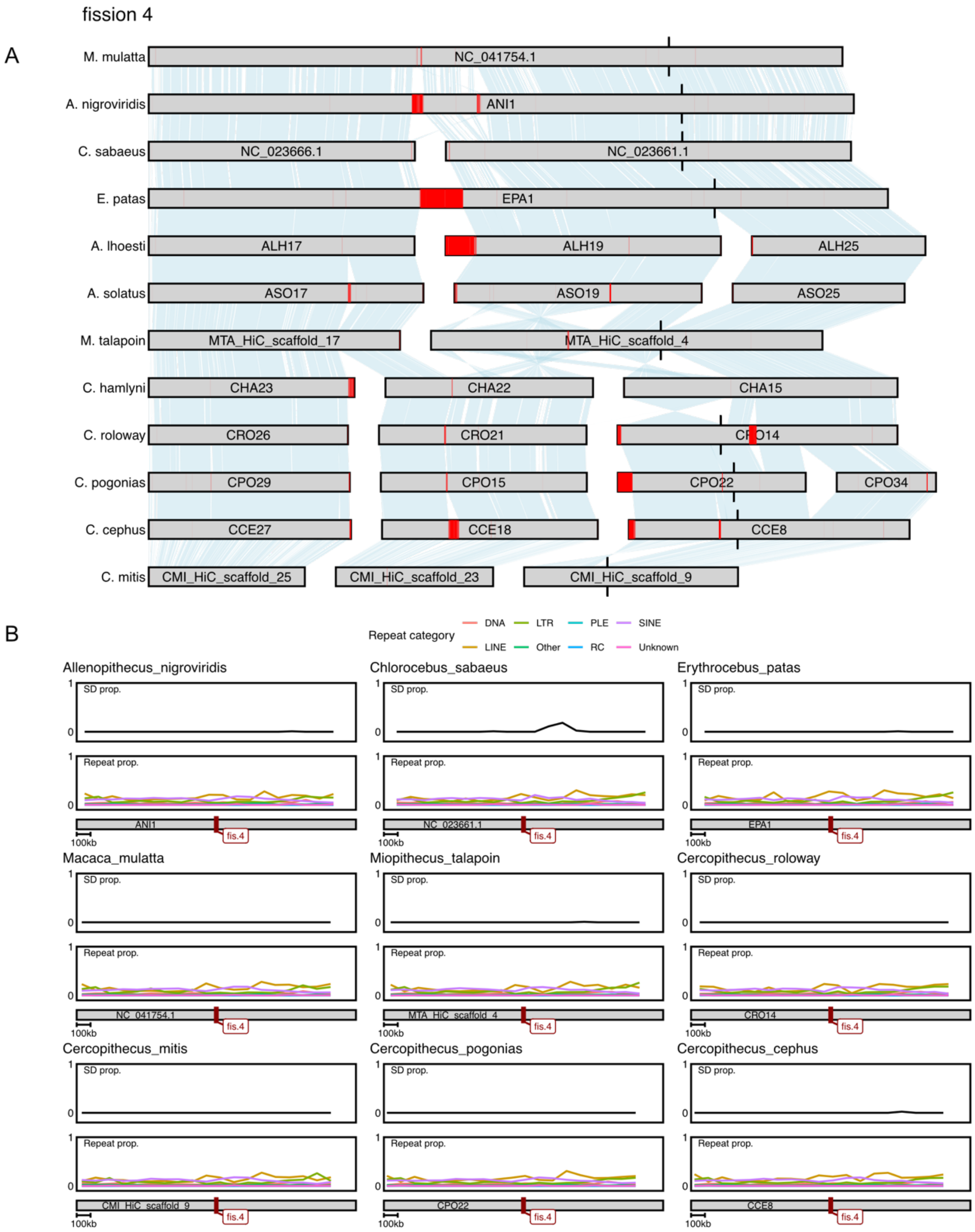


**Figure S12.** A) Chromosomal synteny around the ancestral breaking point of fission 4. Black markings highlight the fission breakpoint, and centromeric annotations from TRASH is highlighted in red. B) Proportion of segmental duplications (SD, top panels) and repetitive elements as annotated by EarlGrey (bottom panels) summarized in 100 kb sliding windows, in lineages lacking the fission. Fission 4 does not overlap putative centromeres in any lineage (A), suggesting this is a non-centromeric fission. No peaks of SD or other repeats are present in outgroup lineages (B).

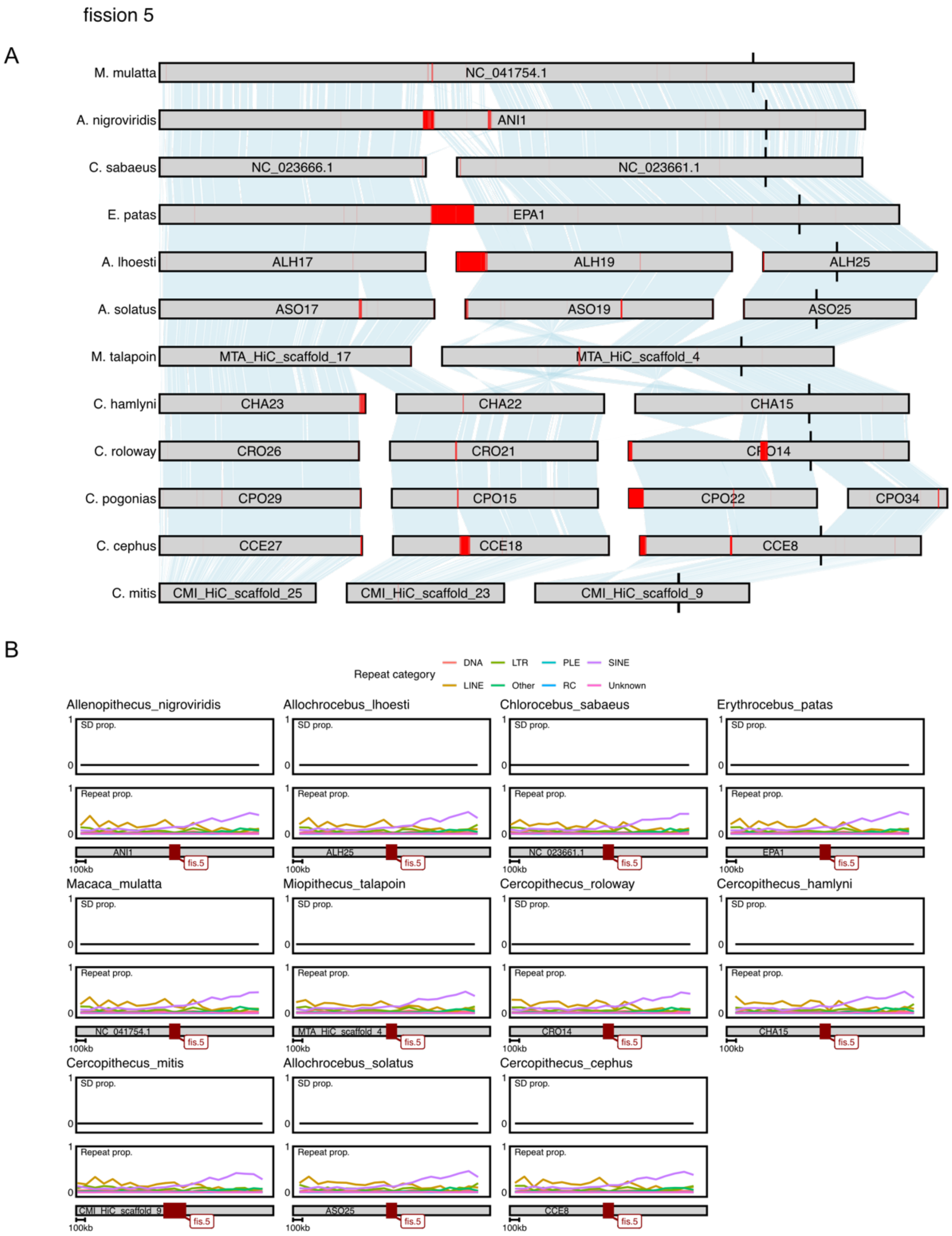


**Figure S13.** A) Chromosomal synteny around the ancestral breaking point of fission 5 Black markings highlight the fission breakpoint, and centromeric annotations from TRASH is highlighted in red. B) Proportion of segmental duplications (SD, top panels) and repetitive elements as annotated by EarlGrey (bottom panels) summarized in 100 kb sliding windows, in lineages lacking the fission. Fission 5 does not overlap putative centromeres in any lineage (A), suggesting this is a non-centromeric fission. No peaks of SD or other repeats are present in outgroup lineages (B).

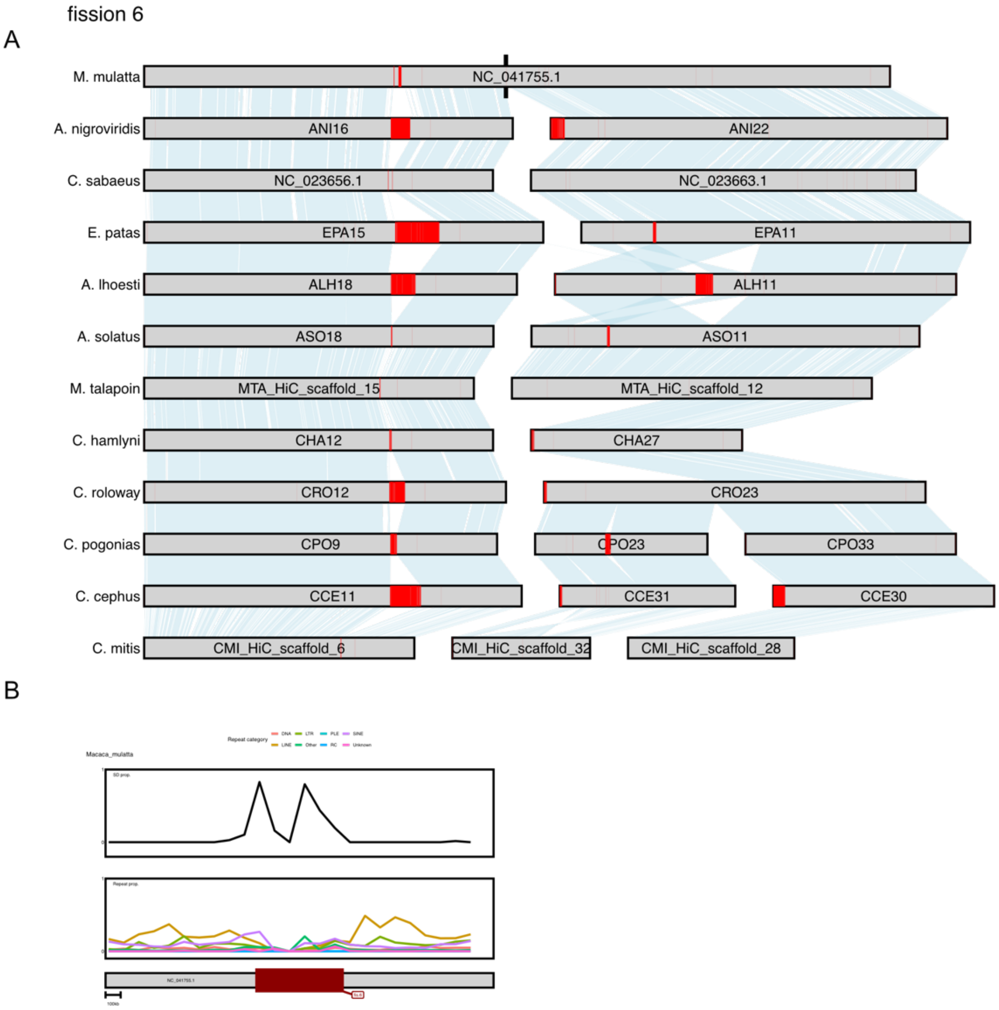


**Figure S14.** A) Chromosomal synteny around the ancestral breaking point of fission 6. Black markings highlight the fission breakpoint, and centromeric annotations from TRASH is highlighted in red. B) Proportion of segmental duplications (SD, top panels) and repetitive elements as annotated by EarlGrey (bottom panels) summarized in 100 kb sliding windows, in lineages lacking the fission. Fission 6 does not overlap putative centromeres in any lineage (A), suggesting this is a non-centromeric fission. The breakpoint overlaps clear peaks of SD content in *M. mulatta* (B).

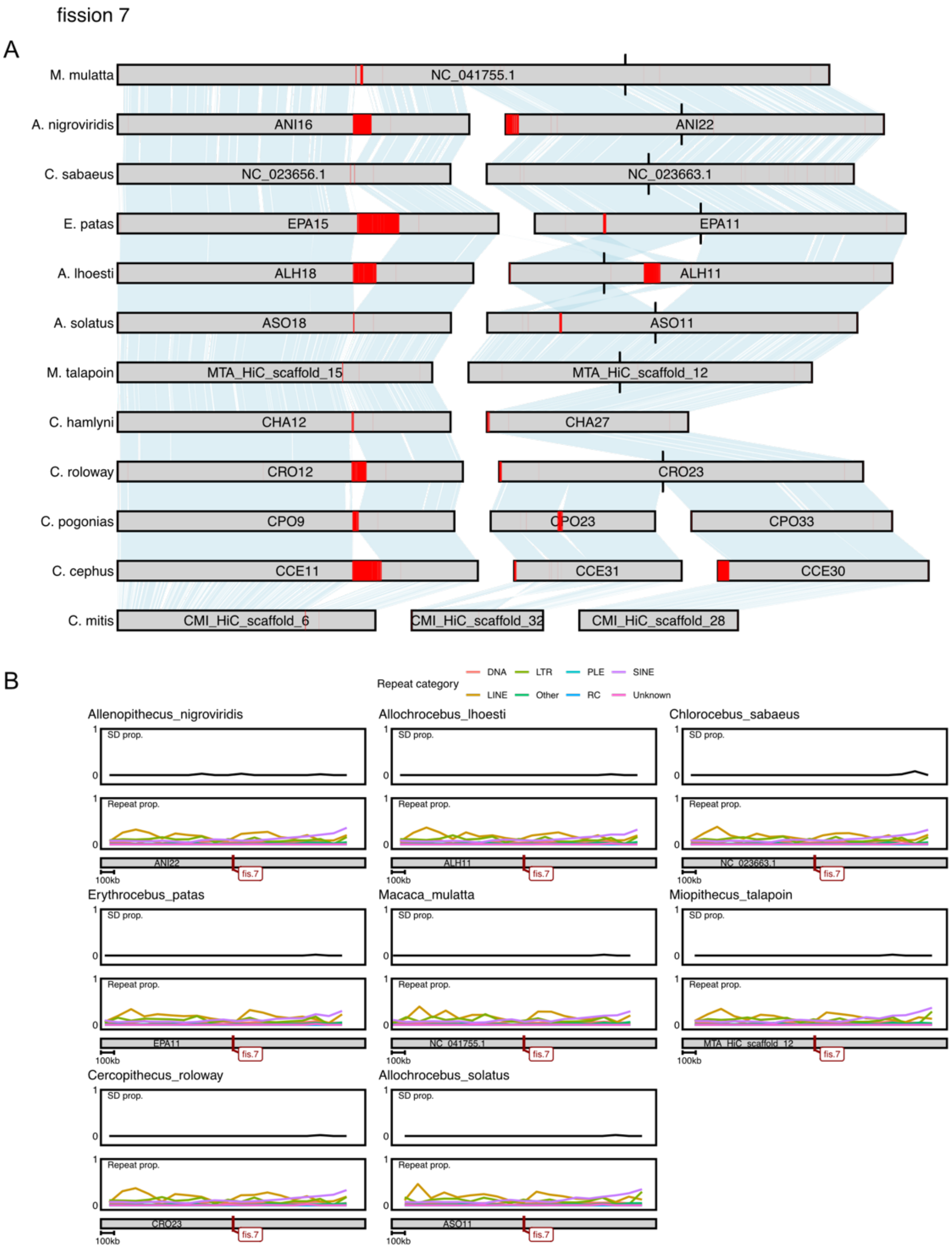


**Figure S15.** A) Chromosomal synteny around the ancestral breaking point of fission 7. Black markings highlight the fission breakpoint, and centromeric annotations from TRASH is highlighted in red. B) Proportion of segmental duplications (SD, top panels) and repetitive elements as annotated by EarlGrey (bottom panels) summarized in 100 kb sliding windows, in lineages lacking the fission. Fission 7 does not overlap putative centromeres in any lineage (A), suggesting this is a non-centromeric fission. No peaks of SDs or repetitive elements are present (B).

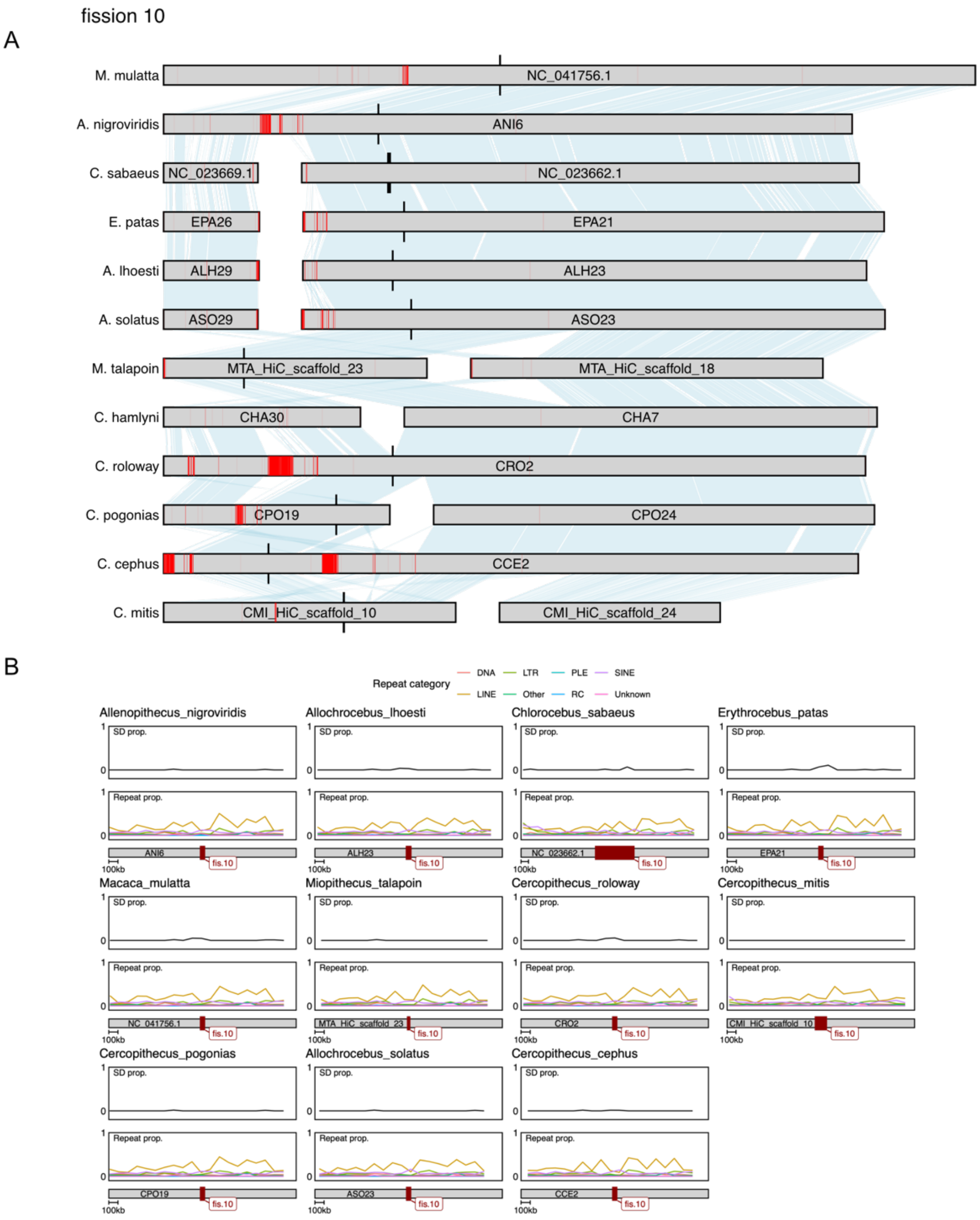


**Figure S16.** A) Chromosomal synteny around the ancestral breaking point of fission 10. Black markings highlight the fission breakpoint, and centromeric annotations from TRASH is highlighted in red. B) Proportion of segmental duplications (SD, top panels) and repetitive elements as annotated by EarlGrey (bottom panels) summarized in 100 kb sliding windows, in lineages lacking the fission. The fission breakpoint does not overlap any putative centromeres in outgroups, and no peaks of SD or other repetitive elements.

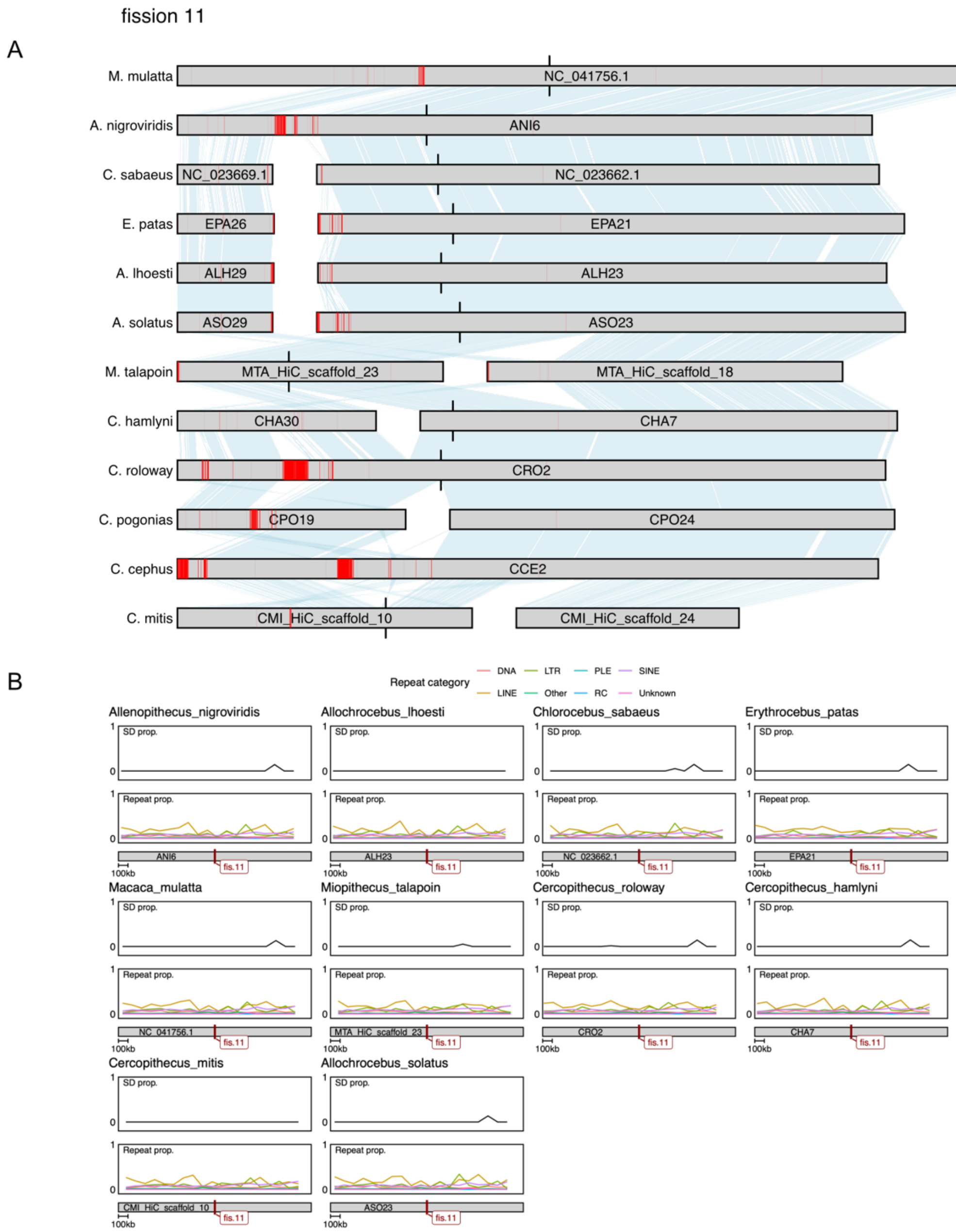


**Figure S17.** A) Chromosomal synteny around the ancestral breaking point of fission 11. Black markings highlight the fission breakpoint, and centromeric annotations from TRASH is highlighted in red. B) Proportion of segmental duplications (SD, top panels) and repetitive elements as annotated by EarlGrey (bottom panels) summarized in 100 kb sliding windows, in lineages lacking the fission. The fission breakpoint does not overlap any putative centromeres in outgroups, and no peaks of SD or other repetitive elements.

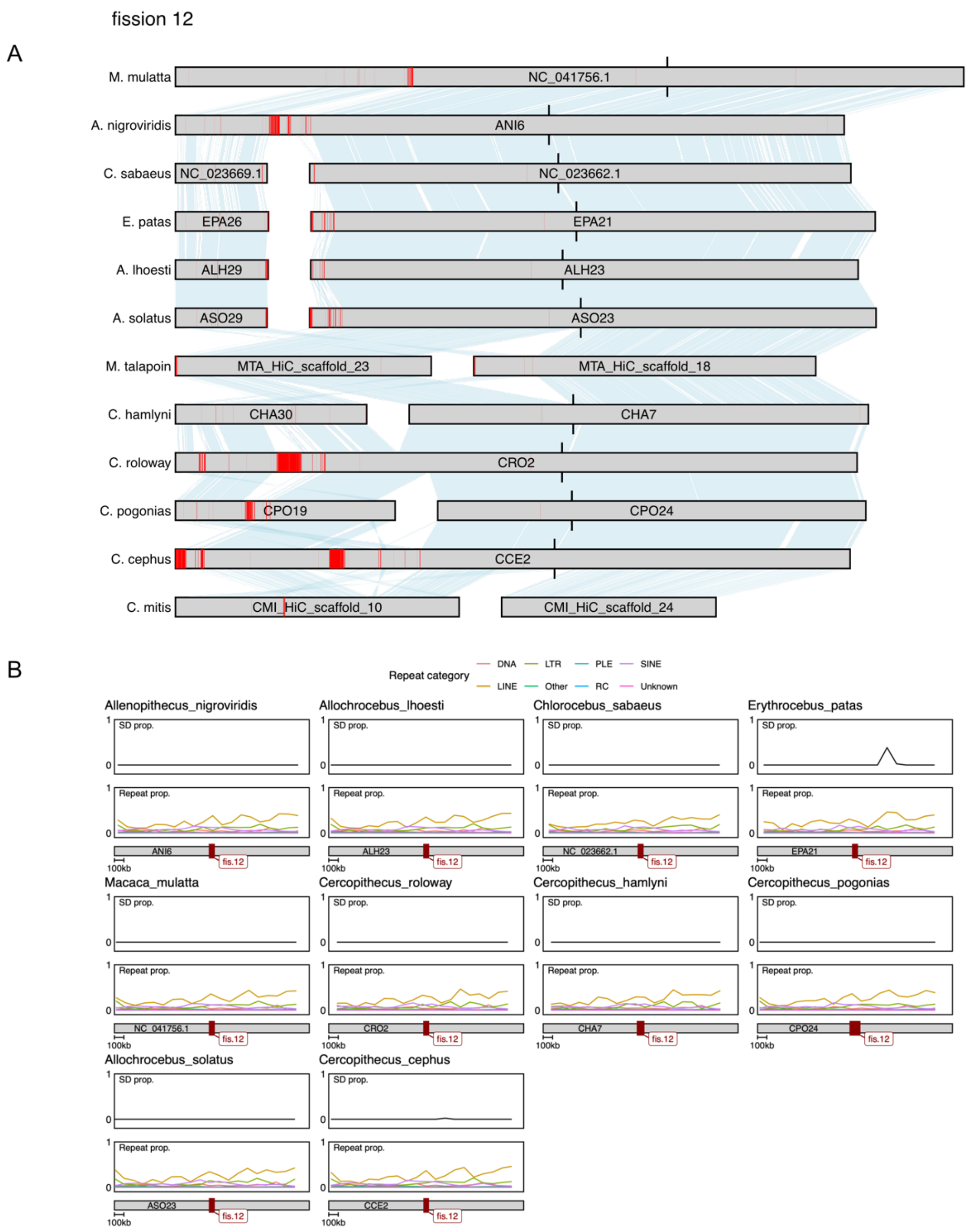


**Figure S18.** A) Chromosomal synteny around the ancestral breaking point of fission 12. Black markings highlight the fission breakpoint, and centromeric annotations from TRASH is highlighted in red. B) Proportion of segmental duplications (SD, top panels) and repetitive elements as annotated by EarlGrey (bottom panels) summarized in 100 kb sliding windows, in lineages lacking the fission. The fission breakpoint does not overlap any putative centromeres in outgroups, and no peaks of SD or other repetitive elements.

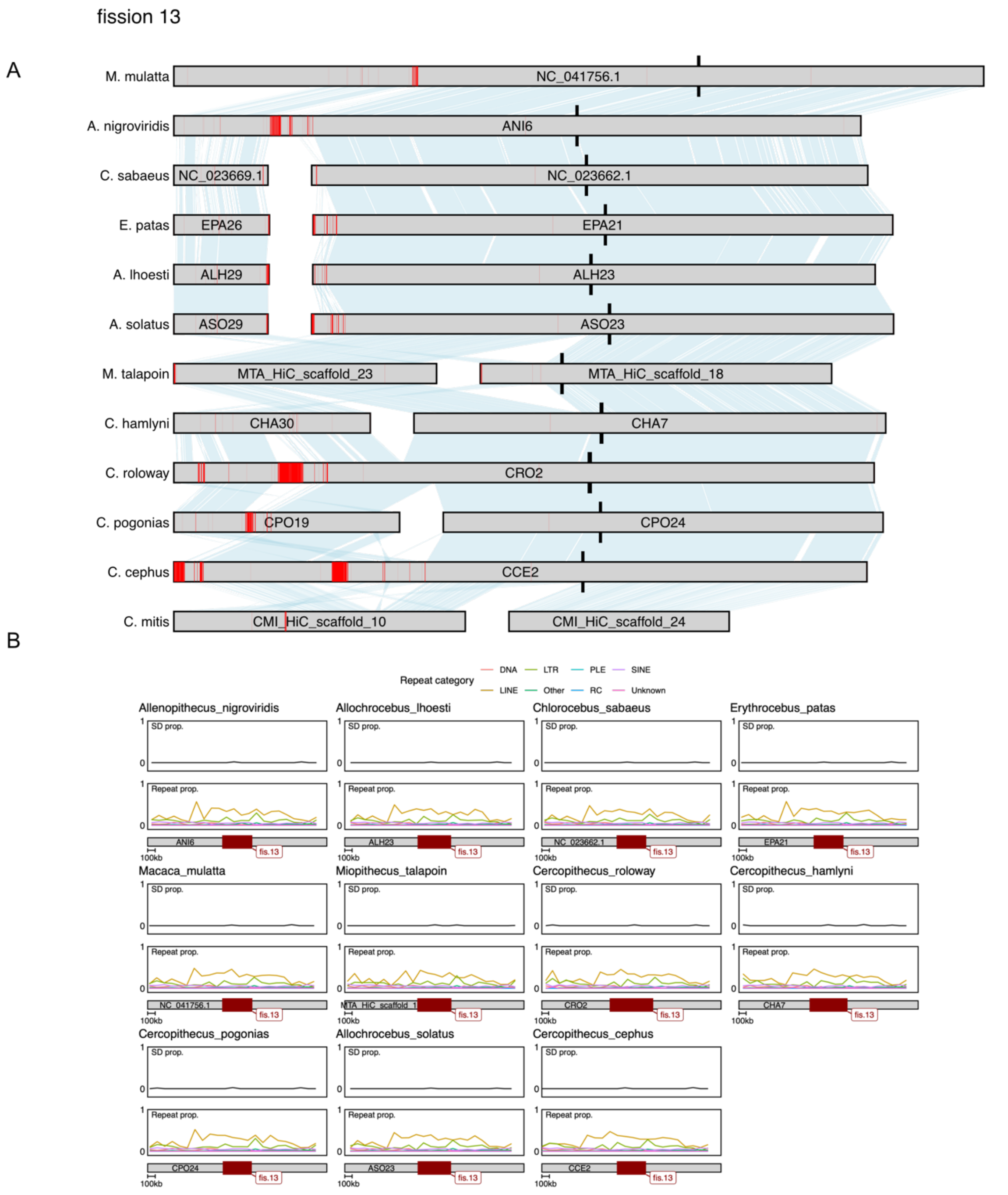


**Figure S19.** A) Chromosomal synteny around the ancestral breaking point of fission 13. Black markings highlight the fission breakpoint, and centromeric annotations from TRASH is highlighted in red. B) Proportion of segmental duplications (SD, top panels) and repetitive elements as annotated by EarlGrey (bottom panels) summarized in 100 kb sliding windows, in lineages lacking the fission. The fission breakpoint does not overlap any putative centromeres in outgroups, and no peaks of SD or other repetitive elements.

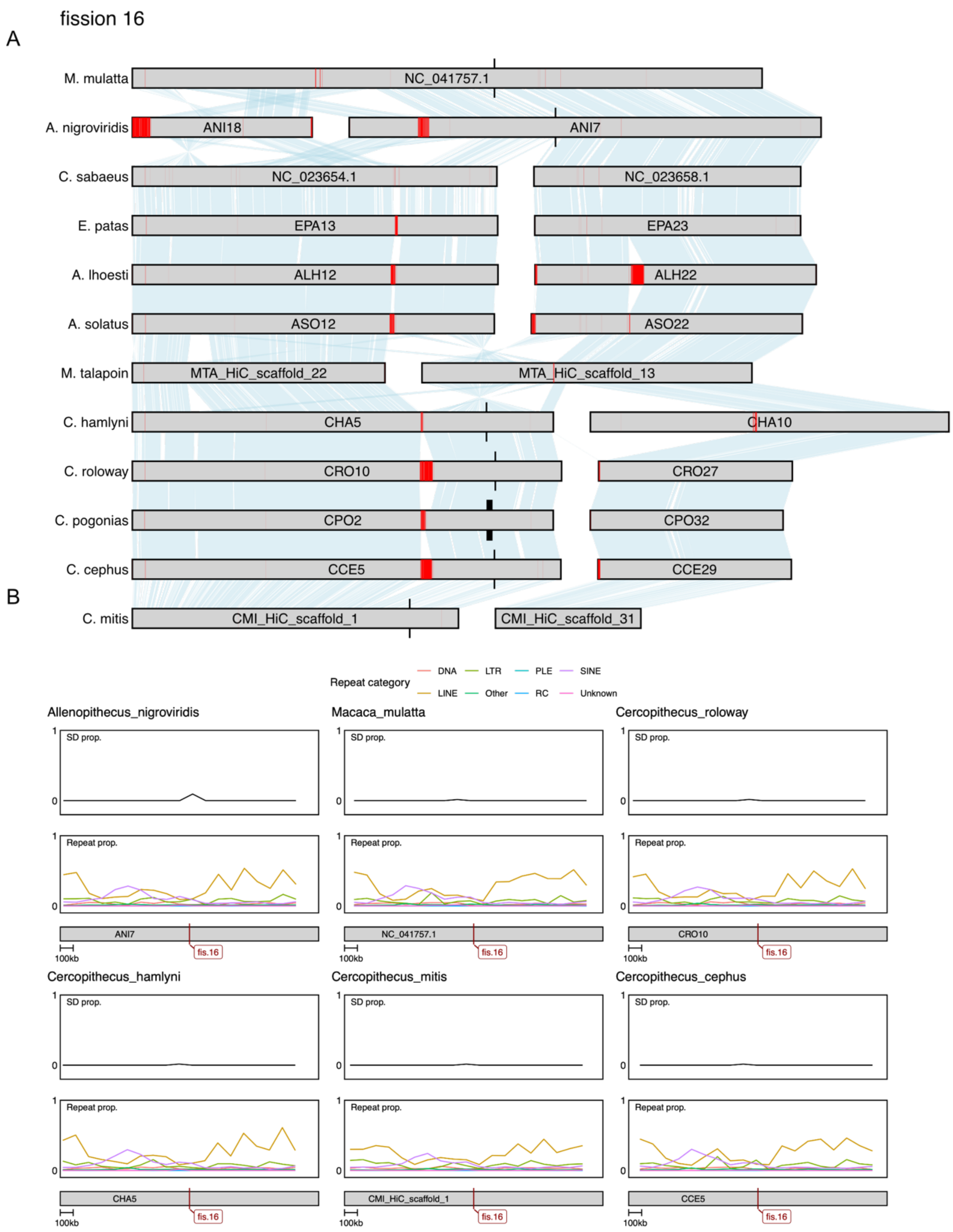


**Figure S20.** A) Chromosomal synteny around the ancestral breaking point of fission 16. Black markings highlight the fission breakpoint, and centromeric annotations from TRASH is highlighted in red. B) Proportion of segmental duplications (SD, top panels) and repetitive elements as annotated by EarlGrey (bottom panels) summarized in 100 kb sliding windows, in lineages lacking the fission. The fission breakpoint does not overlap any putative centromeres in outgroups, and no peaks of SD or other repetitive elements.

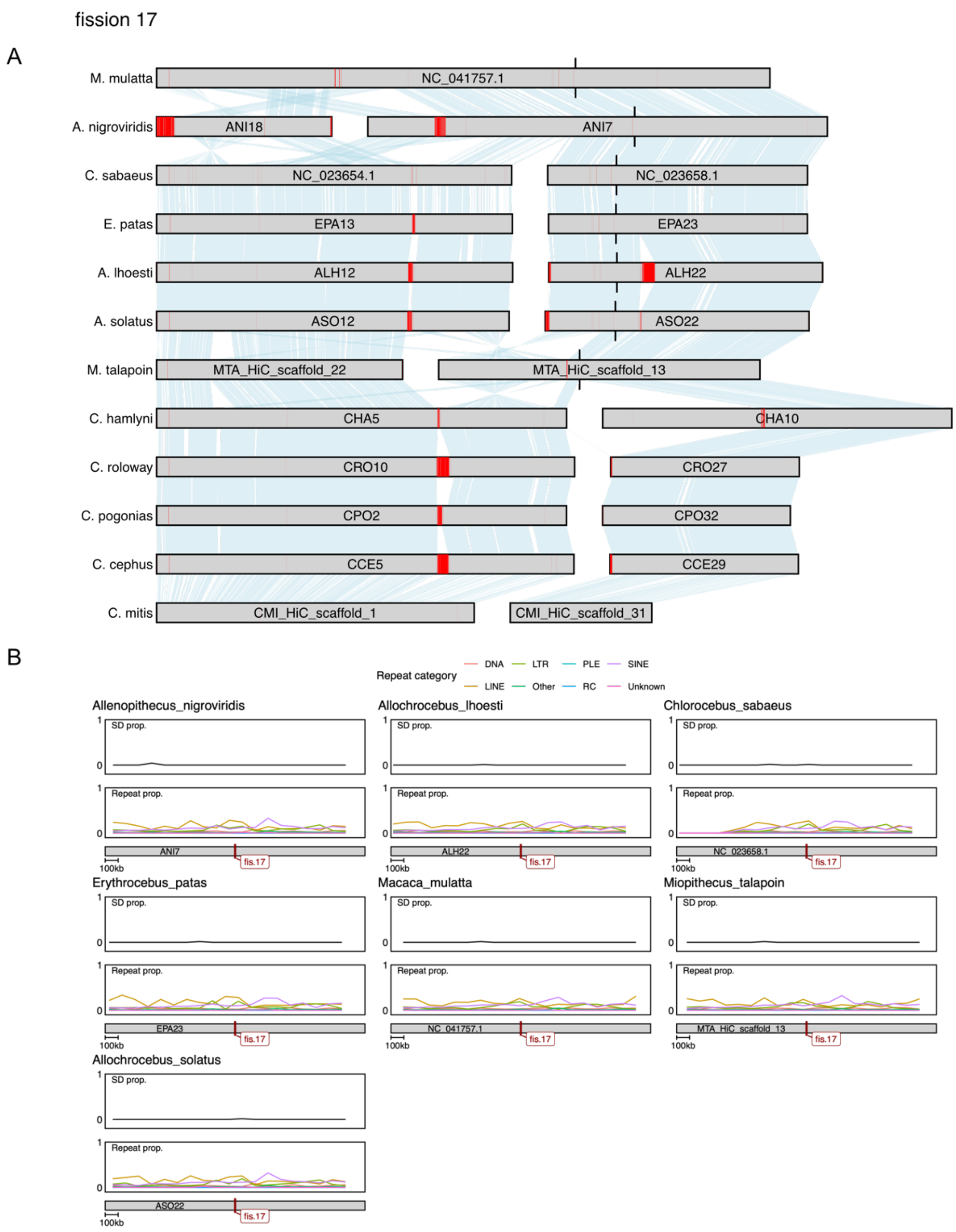


**Figure S21.** A) Chromosomal synteny around the ancestral breaking point of fission 17. Black markings highlight the fission breakpoint, and centromeric annotations from TRASH is highlighted in red. B) Proportion of segmental duplications (SD, top panels) and repetitive elements as annotated by EarlGrey (bottom panels) summarized in 100 kb sliding windows, in lineages lacking the fission. The fission breakpoint does not overlap any putative centromeres in outgroups, and no peaks of SD or other repetitive elements.

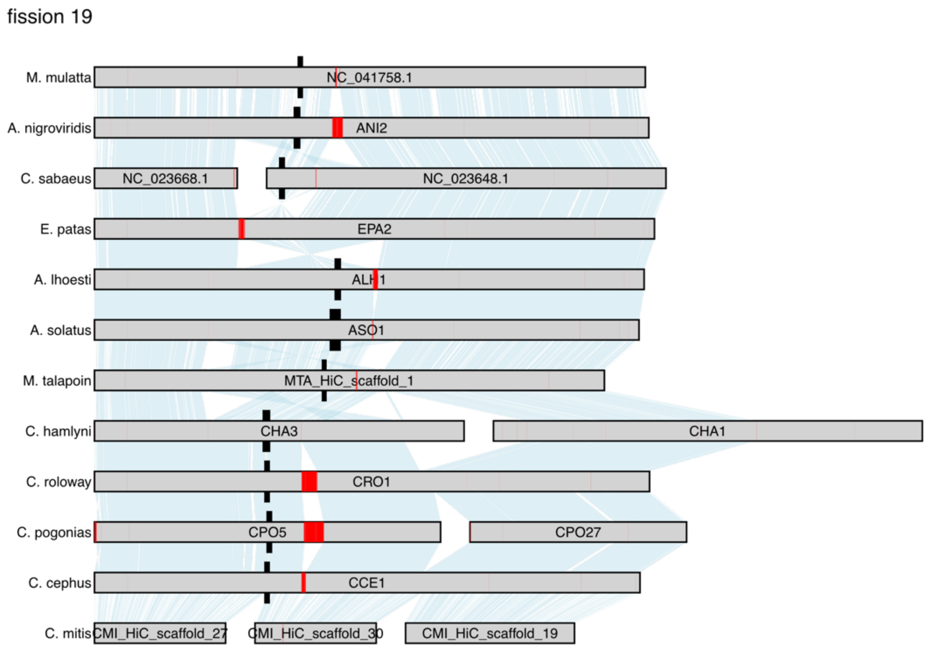


**Figure S22.** Chromosomal synteny around the ancestral breaking point of fission 19. Black markings highlight the fission breakpoint, and centromeric annotations from TRASH is highlighted in red. The fission breakpoint does not overlap putative centromeres in any outgroup, but could not be accurately enough inferred for analyses of SD and repeat content.

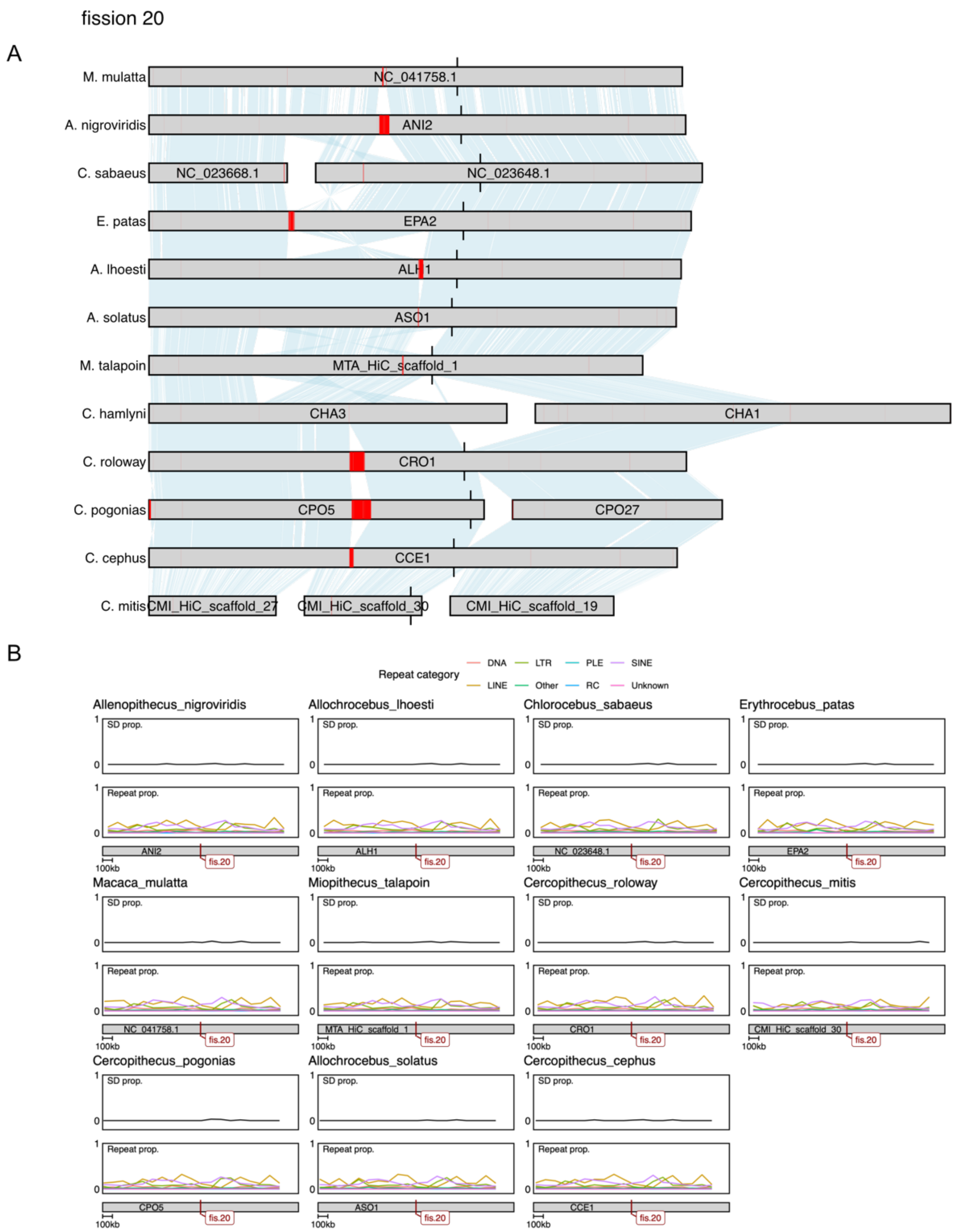


**Figure S23.** A) Chromosomal synteny around the ancestral breaking point of fission 20. Black markings highlight the fission breakpoint, and centromeric annotations from TRASH is highlighted in red. B) Proportion of segmental duplications (SD, top panels) and repetitive elements as annotated by EarlGrey (bottom panels) summarized in 100 kb sliding windows, in lineages lacking the fission. The fission breakpoint does not overlap any putative centromeres in outgroups, and no peaks of SD or other repetitive elements.

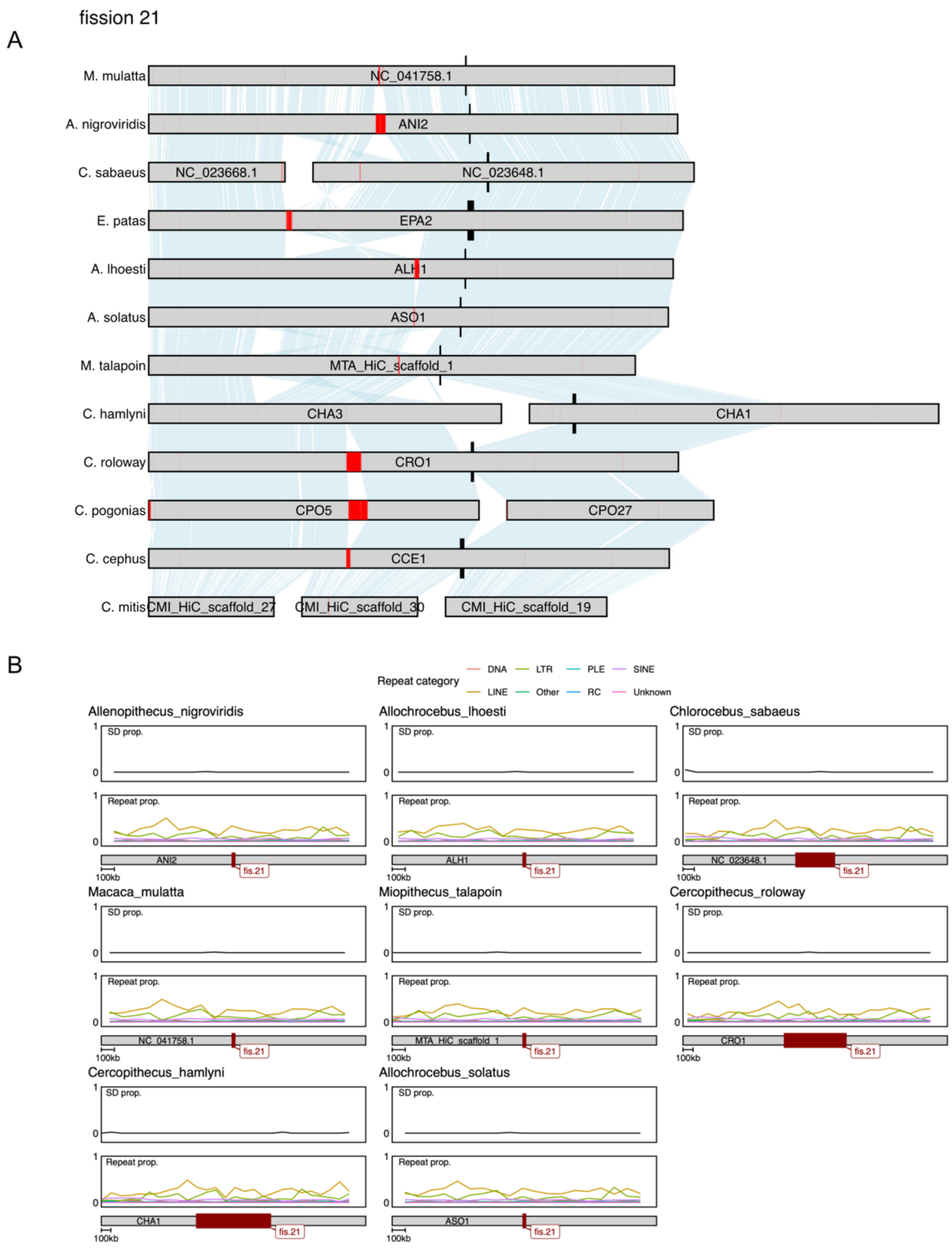


**Figure S24.** A) Chromosomal synteny around the ancestral breaking point of fission 21. Black markings highlight the fission breakpoint, and centromeric annotations from TRASH is highlighted in red. B) Proportion of segmental duplications (SD, top panels) and repetitive elements as annotated by EarlGrey (bottom panels) summarized in 100 kb sliding windows, in lineages lacking the fission. The fission breakpoint does not overlap any putative centromeres in outgroups, and no peaks of SD or other repetitive elements.

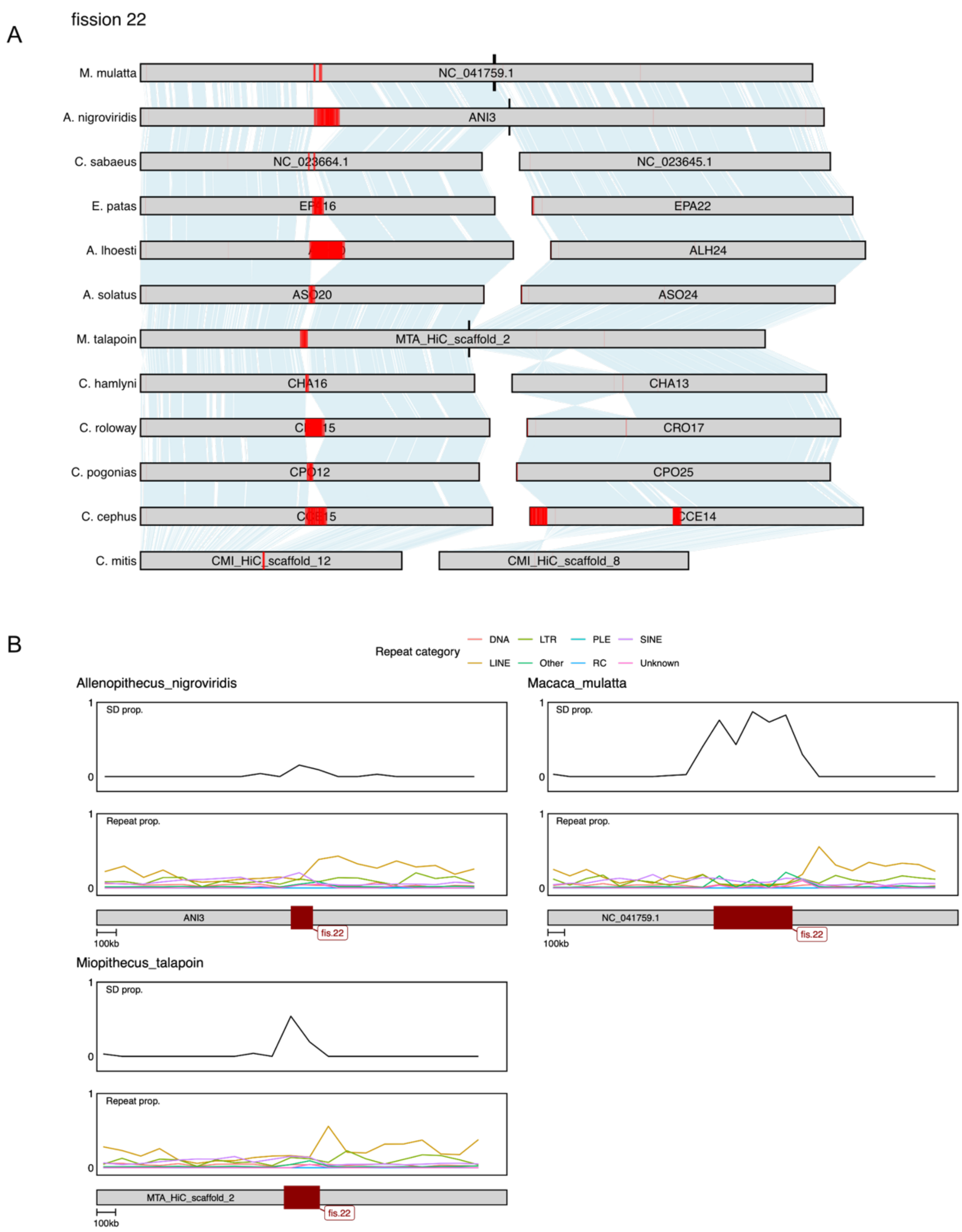


**Figure S25.** A) Chromosomal synteny around the ancestral breaking point of fission 22. Black markings highlight the fission breakpoint, and centromeric annotations from TRASH is highlighted in red. B) Proportion of segmental duplications (SD, top panels) and repetitive elements as annotated by EarlGrey (bottom panels) summarized in 100 kb sliding windows, in lineages lacking the fission. The fission breakpoint does not overlap any putative centromeres in outgroups, but a varying degree of SD peaks are seen in all three species without the fission.

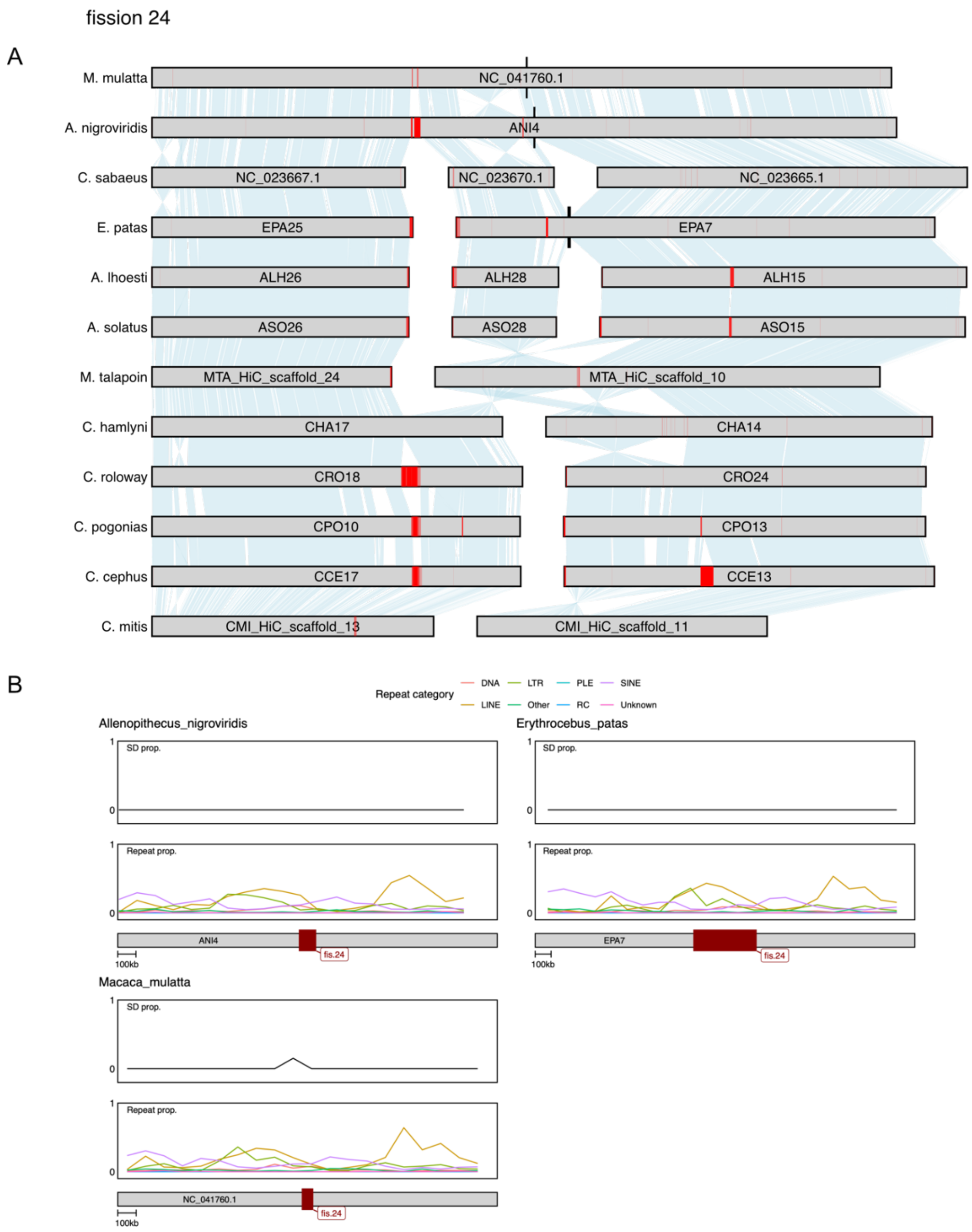


**Figure S26.** A) Chromosomal synteny around the ancestral breaking point of fission 24. Black markings highlight the fission breakpoint, and centromeric annotations from TRASH is highlighted in red. B) Proportion of segmental duplications (SD, top panels) and repetitive elements as annotated by EarlGrey (bottom panels) summarized in 100 kb sliding windows, in lineages lacking the fission. The fission breakpoint does not overlap any putative centromeres in outgroups, and no peaks of SD or other repetitive elements.

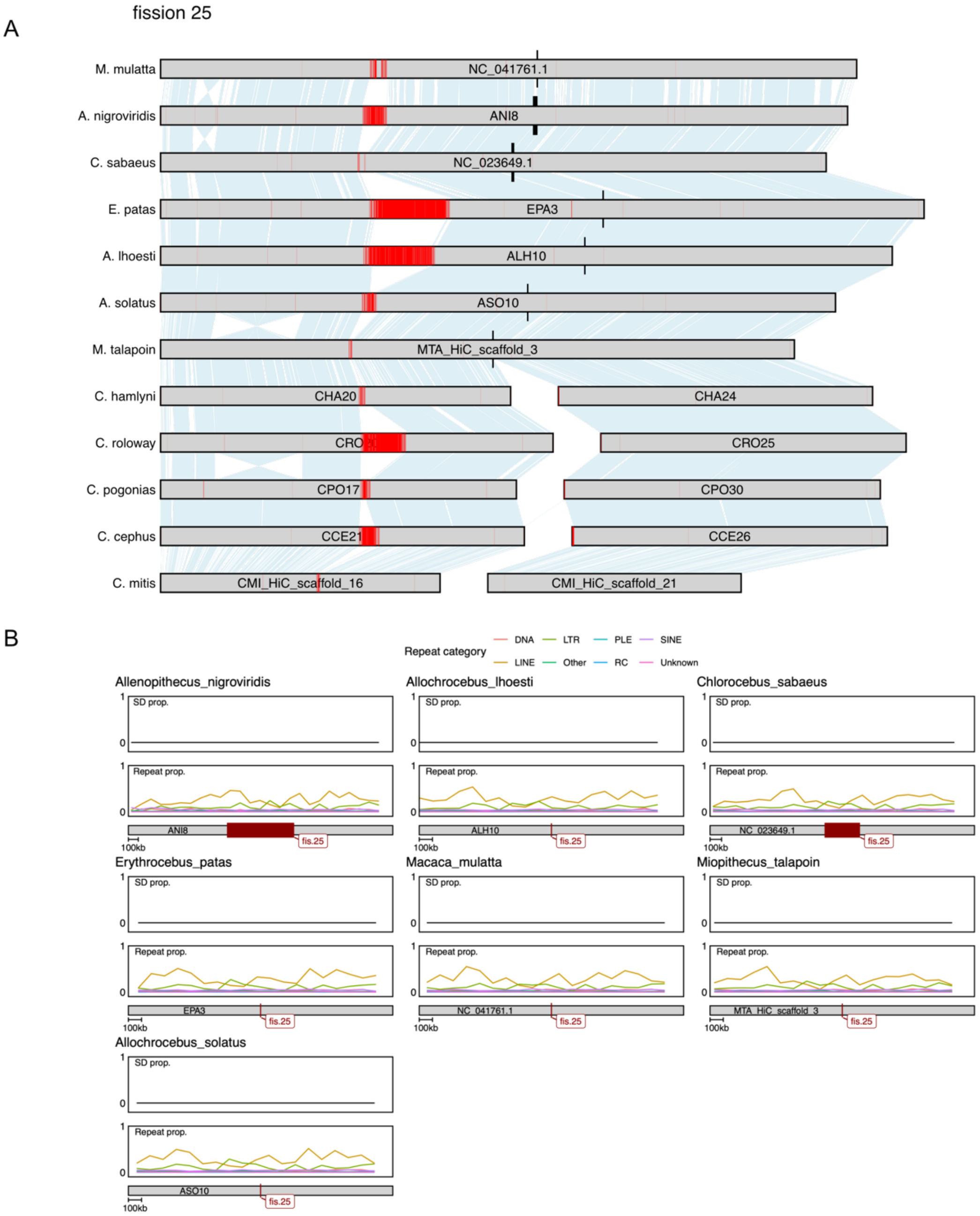


**Figure S27.** A) Chromosomal synteny around the ancestral breaking point of fission 25. Black markings highlight the fission breakpoint, and centromeric annotations from TRASH is highlighted in red. B) Proportion of segmental duplications (SD, top panels) and repetitive elements as annotated by EarlGrey (bottom panels) summarized in 100 kb sliding windows, in lineages lacking the fission. The fission breakpoint does not overlap any putative centromeres in outgroups, and no peaks of SD or other repetitive elements.

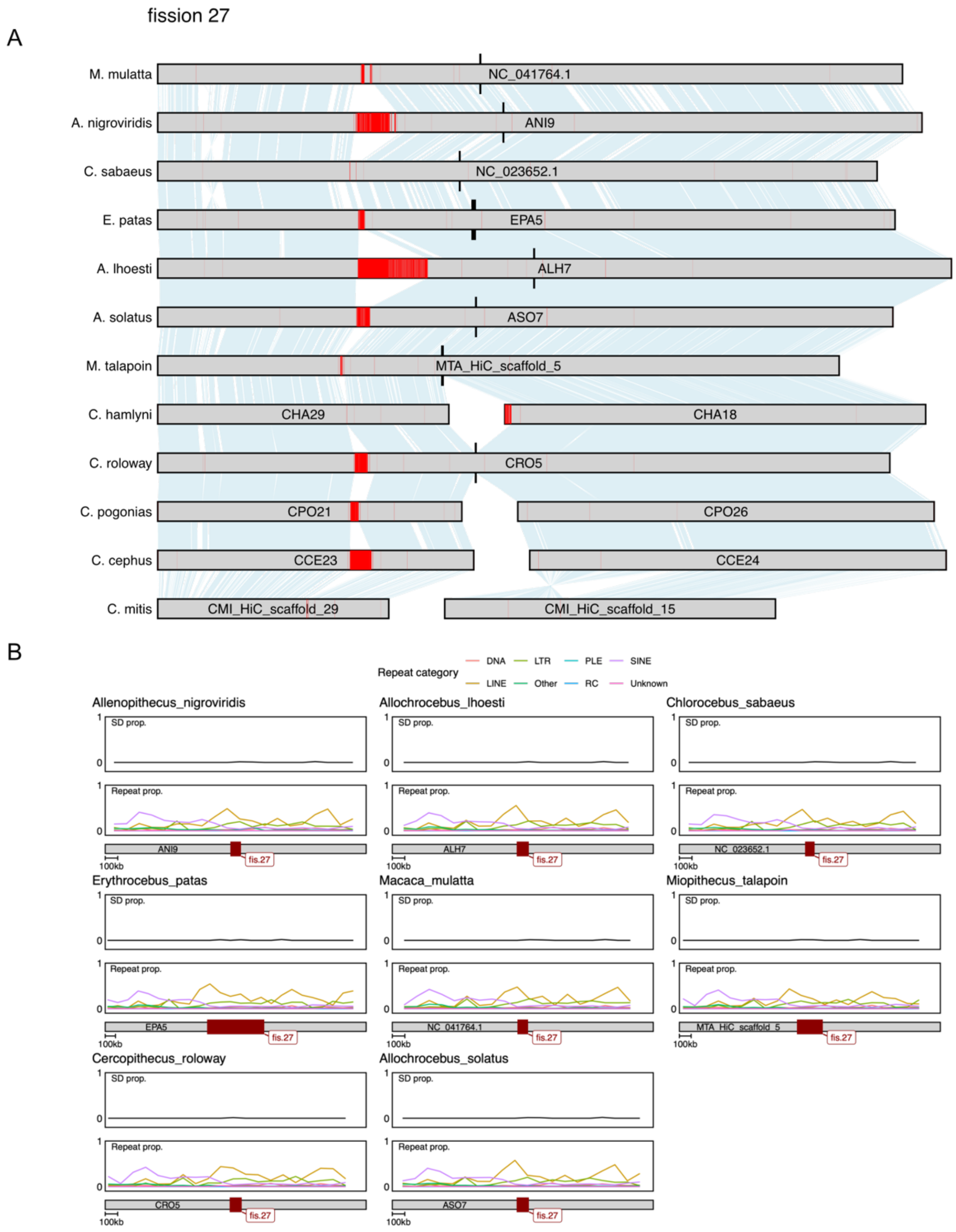


**Figure S28.** A) Chromosomal synteny around the ancestral breaking point of fission 27. Black markings highlight the fission breakpoint, and centromeric annotations from TRASH is highlighted in red. B) Proportion of segmental duplications (SD, top panels) and repetitive elements as annotated by EarlGrey (bottom panels) summarized in 100 kb sliding windows, in lineages lacking the fission. The fission breakpoint does not overlap any putative centromeres in outgroups, and no peaks of SD or other repetitive elements.

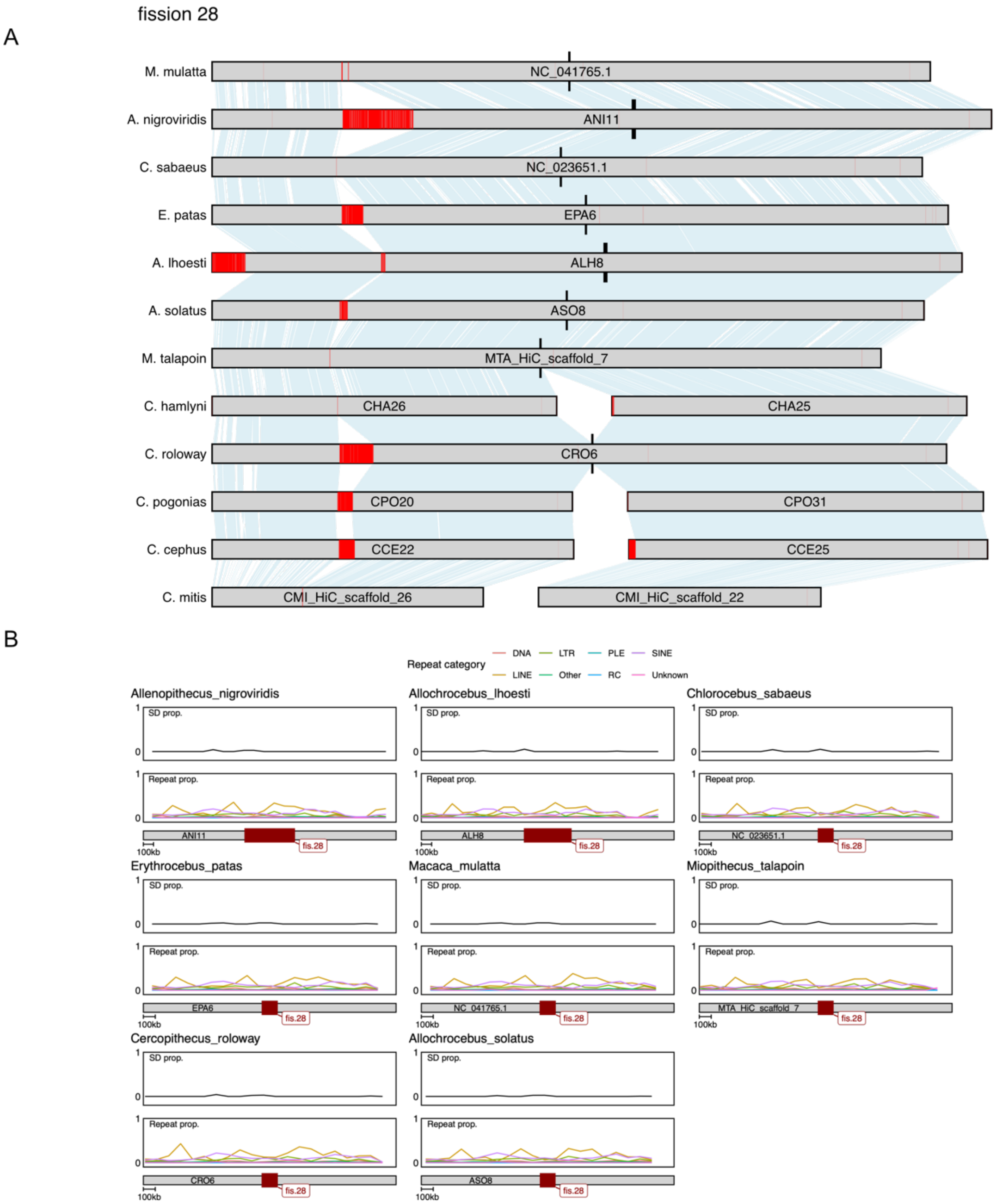


**Figure S29.** A) Chromosomal synteny around the ancestral breaking point of fission 28. Black markings highlight the fission breakpoint, and centromeric annotations from TRASH is highlighted in red. B) Proportion of segmental duplications (SD, top panels) and repetitive elements as annotated by EarlGrey (bottom panels) summarized in 100 kb sliding windows, in lineages lacking the fission. The fission breakpoint does not overlap any putative centromeres in outgroups, and no peaks of SD or other repetitive elements.

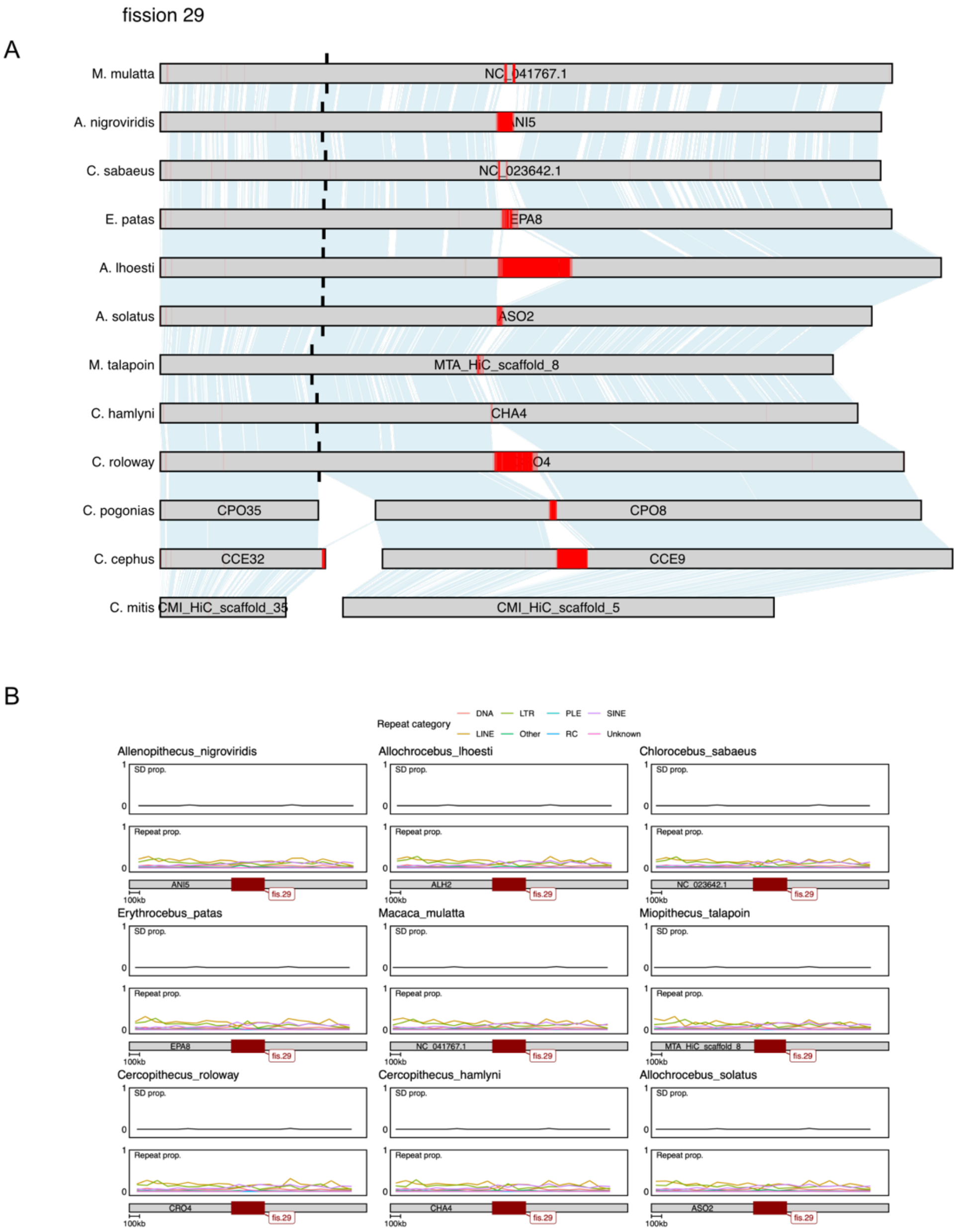


**Figure S30.** A) Chromosomal synteny around the ancestral breaking point of fission 29. Black markings highlight the fission breakpoint, and centromeric annotations from TRASH is highlighted in red. B) Proportion of segmental duplications (SD, top panels) and repetitive elements as annotated by EarlGrey (bottom panels) summarized in 100 kb sliding windows, in lineages lacking the fission. The fission breakpoint does not overlap any putative centromeres in outgroups, and no peaks of SD or other repetitive elements.
